# A neural signature for the subjective experience of threat anticipation under uncertainty

**DOI:** 10.1101/2023.09.20.558716

**Authors:** Xiqin Liu, Guojuan Jiao, Feng Zhou, Keith M Kendrick, Dezhong Yao, Shitong Xiang, Tianye Jia, Xiaoyong Zhang, Jie Zhang, Jianfeng Feng, Benjamin Becker

## Abstract

Uncertainty about potential future threats and the associated anxious anticipation represents a key feature of anxiety. However, the neural systems that underlie the subjective experience of threat anticipation under uncertainty remain unclear. Combining a novel uncertain shock anticipation paradigm that allows precise modulation of the level of momentary anxious arousal during functional magnetic resonance imaging (fMRI) with multivariate predictive modeling, we trained a brain model that accurately predicted the intensity of subjective experience of anxious arousal on the population and individual level. In a series of analyses utilizing available fMRI datasets, we further demonstrate that the signature specifically predicted anxious anticipation and was not sensitive in predicting pain, general anticipation or unspecific arousal. The signature was functionally and spatially distinguishable from representations of subjective fear or negative affect. We developed a sensitive, generalizable, and specific neuromarker for subjective anxious arousal experienced during uncertain threat anticipation that can facilitate model development and clinical translation.

## Introduction

Uncertainty refers to the inability to predict the outcome of a situation, or the likelihood, valence, intensity, time, or type of future events^1^. It represents an essential part of our daily lives ranging from uncertainty in the early stages of a romantic relationship to the likelihood of catching a COVID-19 infection or the magnitude of climate change. Uncertainty about potential threats in the future and the associated anticipatory processes are central to the feeling of anxiety^2,3^. While anxiety serves an important adaptive function to avoid or cope with potential danger^4^, excessive anticipatory anxiety in the face of uncertainty represents a key symptom of anxiety disorders^2,5^. Contemporary neurobiological frameworks for mental disorders therefore consider ‘potential threat’ and ‘uncertainty intolerance’ as candidate mechanisms of anxiety (e.g., the Research Domain Criteria, RDoC)^6,7^. However, the neural pathways that underlie the actual subjective experience of uncertainty-induced threat anticipation remain unclear.

Uncertain threat anticipation represents a prototypical paradigm to evoke experimental anxiety^2,8,9^. Rodent models have identified brain systems that mediate the behavioral and physiological responses to uncertain situations (e.g., shock-probe burying)^10^. The bed nucleus of the stria terminalis (BNST), for instance, critically mediates defensive behaviors (e.g., freezing, flight and avoidance) during uncertain threat anticipation in rodents^11–14^ and has also been proposed as a key anxiety system in the RDoC framework^6,7^. Other core regions include the medial prefrontal cortex (mPFC), ventral hippocampus (vHPC), amygdala, insula and thalamus^10,13,15^. Recent technological advances (e.g., optogenetics) in animal research have reconciled the region-focused perspective into circuit-level frameworks demonstrating that anxiety-related responses are mediated by distributed circuits^10^. However, the contribution of these neural mechanisms to the conscious experience of anxiety remains unknown because subjective feelings cannot be assessed in animal models^16,17^. Circuits that underlies defensive behaviors are also distinct from those that generate subjective emotional experiences^18–22^. Given current treatments based on behavioral and physiological indices are less effective than desirable^23,24^, and feelings of excessive anxiety are the primary reason for patients to seek treatment and reduction of subjective symptoms represents treatment success^22^, a mechanistic understanding of the neural representation that supports the subjective experience of uncertain threat anticipation can facilitate the development of clinical biomarkers and new treatments for anxiety^10,18,22,25,26^.

Human functional magnetic resonance imaging (fMRI) studies have reported BNST activation to uncertain threat anticipation^27–32^, yet evidence that the BNST encodes subjective anticipatory experience remains controversial. Previous fMRI studies comparing conditions of uncertain threat versus safe anticipation (overview see^33^) have revealed increased activity in a broad range of brain regions such as BNST^30,31,34,35^, amygdala^36–38^, periaqueductal gray (PAG)^29,34,39^, anterior insula (aINS)^27–29,40,41^, anterior cingulate cortex (ACC)^28,29,40,42,43^ and lateral and medial frontal regions^41,44–46^. However, the comparison does not allow to specifically isolate the subjective feeling of uncertain threat anticipation given that the conditions may differ in several other mental processes (e.g., defensive responses or arousal), and the identified regions are involved in fundamental cognitive processes including salience or arousal^13^. Emotional experiences are highly subjective yet accessible via introspective self-report^47^. Recent constructionist theories suggest that the subjective experience is supported by a distributed set of interacting brain regions involved in emotional and non-emotional operations^48,49^. However, there is limited research focusing on the brain mechanism of actual subjective experiences during uncertain threat anticipation and methodological limitations of the prevailing neuroimaging designs and analytic approaches makes it difficult to isolate its specific neural substrates^21,22^.

One fMRI study determined the neural basis of subjective experience of threat anticipation as a function of trial-by-trial self-reported level of anxious arousal during aversive and neutral auditory trials and showed an association between subjective anxious feelings and reactivity in amygdalo-insular systems^50^. However, a conventional local mapping (mass-univariate) approach was employed in this study to identify isolated brain regions associated with subjective ratings. This approach tests hypotheses about structure–function associations and lacks functional specificity to provide a sufficient brain-level model of an emotional state^51^. A growing body of research suggests that many cognitive and emotional processes involve distributed neural coding across multiple brain regions or networks^52–54^, and it has been proposed to examine population coding of neural activity in terms of multivariate activation patterns^55^. Multivariate pattern analysis (MVPA), therefore, has emerged as a powerful method for capturing emotion-specific brain states at a fine-grained level^56^. Specifically, multivariate predictive modeling, a machine learning technique based on MVPA, is suggested to be more suitable for identifying neuromarkers for specific subjective emotional states with high effect sizes and clinical significance by providing predictions (instead of local mapping) about emotional experience from distributed neural activity patterns^51,57^. Chang et al. showed for instance that a whole-brain multivariate model explained much more variance in predicting experiencing negative affect than local regions^51,52^, and this predictive modeling approach has been successfully utilized to develop models that can sensitively and specifically predict subjective experiences of fear^21^,pain^53^ and pleasure^58^.

Here, we developed an ‘uncertainty-variation threat anticipation (UVTA)’ fMRI paradigm that modulates different aspects of shock uncertainty to capture momentary variations in subjective reports of anxious arousal. Using multivariate predictive modeling, we aimed to determine (1) whether it is possible to develop a process-specific, robust and generalizable neural representation (‘signature’) predictive of the intensity of subjective anxious experience during uncertain threat anticipation on the population level, (2) whether this signature can accurately track momentary trial-by-trial variations of anxious anticipation on the individual level, and (3) whether and which regions make consistent contributions to the whole-brain predictive models. Next, we systematically determined the extent to which the signature (4) depends on unspecific processes (e.g., negative emotional and autonomic arousal, anticipation per se), (5) differs from signatures of subjective fear or general negative experience, and (6) whether regions such as BNST or ‘salience network’ are sufficient to predict subjective anxious arousal.

To this end, we used 9 datasets (Studies 1–11, *n* = 572), including three new experiments (Studies 1–3, *n* = 124) during which participants experienced varying levels of anticipatory anxious arousal induced by the UVTA paradigm (**Fig. 1a**). The UVTA paradigm was based on the established ‘threat of shock (TOS)’ paradigm frequently used to induce anxiety during the anticipation of uncertain electric shocks in experimental contexts (see ref.^9^). We here modulated uncertainty along different dimensions (occurrence, timing and number of shocks) to induce within-individual variations in subjective ratings of anxious arousal (**Fig. 1b**). We trained a predictive model of ratings using support vector regression (SVR) on whole-brain activity patterns (Study 1, *n* = 44), termed the ‘shock uncertainty-induced threat anticipation signature’ (SUITAS), which was evaluated in a validation dataset (Study 2, *n* = 30, identical paradigm, **Fig. 1c**). Using a prospective independent dataset (Study 3, *n* = 50, modified paradigm) and two publicly available datasets (Study 4–5, *n* = 127), we tested the generalizability of SUITAS across cohorts, paradigms, MRI systems and scanning parameters. Next, four independent datasets were used to demonstrate the specificity of the SUITAS for subjective anxious arousal rather than pain experience (Study 6, *n* = 33), general anticipation (Study 7, *n* = 100), unspecific negative emotional (Study 8, *n* = 48, subsample of Study 3) and autonomic arousal (Study 9, *n* = 65, subsamples of Studies 2 and 3; **Fig. 1d**). Finally, two additional datasets were used to determine whether the SUITAS is functionally and topographically distinguishable from established signatures for subjective fear (Study 10, *n* = 67)^21^ and subjective negative affect (Study 11, *n* = 121)^52^ (**Fig. 1d**). This design allowed a sufficient brain-level description of subjective experience of threat anticipation under uncertainty and we hypothesized that the subjective anxious experience would be decoded by a sensitive, generalizable and specific distributed neural representation that is (partly) separable from representations of fear exposure and negative affect.

**Fig. 1.**
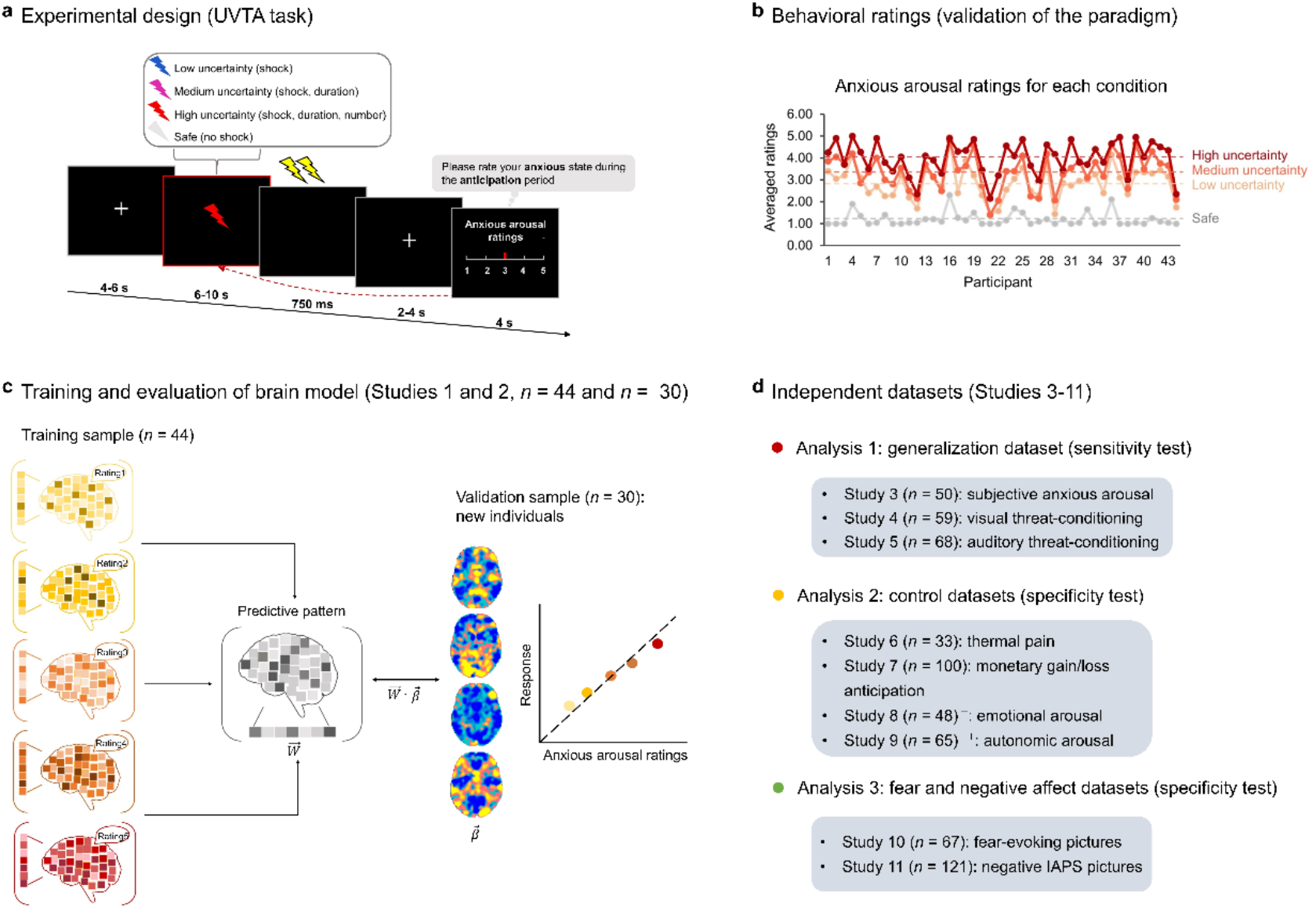
Experimental design and analytic strategy. **a**, Participants anticipated a highly aversive outcome (shock) during the cue periods. Four cue types signaled four uncertainty levels for a total of 96 randomized trials distributed over four fMRI runs. Each trial began with a 4∼6 s fixation-cross followed by a fixed 8 s (blue) or 6∼10 s (avg 8 s) anticipatory cue (purple, red and white) period. When the anticipatory cue disappeared, an outcome (either 2∼3 shocks or no shocks depending on the color of the cue) was delivered to the participants and a black screen was presented for 750 ms. After a 2∼4 s fixation cross, participants reported their level of anxious arousal during the preceding anticipatory cue presentation phase on a Likert scale ranging from 1 to 5 (1 = no anxiety to 5 = extreme anxiety) within 4 s. **b**, Averaged anxious arousal ratings according to the four different uncertainty level conditions plotted for each participant in Study 1 (*n* = 44). The dash lines indicate the mean ratings for each condition. The paradigm robustly induced different levels of anxious arousal, with most participants reporting a broad range of anticipatory arousal levels. **c**, The training of multivariate signature for subjective anxious arousal using a support vector regression model for the anticipation period (6∼10 s, Study 1, *n* = 44) and evaluation of the signature using new individuals (Study 2, *n* = 30). **d**, Validation on independent datasets to test the signature’s generalizability (Study 3, *n* = 50; Study 4, *n* = 59; Study 5, *n* = 68), specificity with respect to neural activity during related processes including pain (Study 6, *n* = 33), positive and negative anticipation of non-shock events (Study 7, *n* = 100), negative emotional arousal (Study 8, *n* = 48), autonomic arousal (Study 9, *n* = 65) and predictive and topological specificity in comparison to subjective fear (Study 10, *n* = 67) and general negative affect (Study 11, *n* = 121). ^+^ The sample (*n* = 48) in Study 8 was a subsample of Study 3 (*n* = 50). ^++^ The sample (*n* = 65) in Study 9 was a subsample of Study 2 (*n* = 30) and Study 3 (*n* = 50).

## Results

### Validation of the UVTA paradigm

In Study 1–3, participants underwent the event-related UVTA paradigm, a modified version of the classical ‘TOS’ anxiety induction paradigm^9^. Participants were instructed to anticipate potential threats (highly aversive electric shocks) with four different uncertainty levels (i.e., certain safety, low, medium and high uncertainty) varied in occurrence, timing and number of shocks indicated by four different cues (Fig. 1a). For the certain safety condition, no shocks would be administered after a white cue (6∼10 s); for low uncertainty condition, two consecutive shocks might occur immediately after a blue cue (8 s); for medium uncertainty condition, two consecutive shocks might occur immediately after a purple cue (6∼10 s); for high uncertainty condition, two or three shocks might occur immediately after a red cue (6∼10 s) (details see Methods). At the end of each trial, participants retrospectively rated their subjective level of anxious arousal during the anticipation period (6∼10 s) from 1 (no anxiety) to 5 (extreme anxiety) (for details, see Fig. 1a and Methods). The paradigm robustly induced the entire range of anxious arousal, such that 84%, 87% and 98% of participants rated ‘1-4’ and 59%, 50% and 80% of participants reported all 5 levels of anxious arousal in Study 1, 2 and 3, respectively.

To validate whether different uncertainty conditions resulted in varying subjective ratings of anxious anticipation, we estimated linear mixed-effects models (LMMs) with self-reported ratings as the dependent variable and uncertainty condition as the independent variable in Study 1, 2 and 3, respectively. We observed main effects of conditions in all three studies (Study 1: *F*_(3,129)_ = 367.00, *P* < 0.001, *η^2^_p_* = 0.90 [0.87,1.00]; Study 2: *F*_(3,87)_ = 164.90, *P* < 0.001, *η^2^_p_* = 0.85 [0.80,1.00]; Study 3: *F*_(3,147)_ = 448.54, *P* < 0.001, *η^2^_p_* = 0.90 [0.88,1.00]), with subjective ratings increased with uncertainty levels (safety < low < medium < high, all post-hoc *P*s < 0.001, Fig. 1b and Supplementary Fig.1), suggesting that our paradigm robustly induced varied levels of anxious arousal.

Moreover, we asked whether participants’ subjective ratings for different uncertainty conditions were affected by personality traits such as trait anxiety (TA, measured by the Spielberger State-Trait Anxiety Inventory, STAI^59^) and intolerance of uncertainty (IOU, measured by the Intolerance of Uncertainty Scale-short form, IUS-12^60^). LMM analyses revealed that higher IOU scores were linked to higher anxious experience in uncertain threat conditions (with safety as baseline), yet no effect of TA scores was observed (see Supplementary Results and Supplementary Fig. 2), confirming a specific involvement of intolerance of uncertainty in the anxiety-provoking processes of our paradigm.

### SUITAS – a sensitive neural signature predictive of shock uncertainty-induced threat anticipation

We applied machine learning-based MVPA techniques using an SVR algorithm (i.e., predictive modeling) to develop a population-level whole-brain signature in the training dataset [Study 1, *n* = 44; 198 beta images total, 3∼5 images per participant, gray matter mask, modeled for entire anticipation period (6∼10 s), see Methods]. Then, we evaluated the performance of the SUITAS by conducting 10 × 10-fold cross-validation and applying the SUITAS to new individuals in the validation dataset (Study 2, *n* = 30) to calculate the SUITAS pattern expressions for each participant in Study 2 (entire anticipation period, one map per rating, Fig. 1c; see also Methods). The SUITAS accurately predicted subjective ratings in both the training and validation datasets and the overall prediction-outcome correlation coefficients were 0.59 (explained variance score (EVS) = 24%; bootstrapped 95% confidence interval (CI) = [0.48, 0.69], *P* < 0.001; within-participant *r* = 0.74 ± 0.01, mean EVS = 45.6 ± 2.3%) and 0.61 (EVS = 28%; bootstrapped 95% CI = [0.50, 0.70], one-tailed permutation test *P* < 0.001; within-participant *r* = 0.77 ± 0.03, mean EVS = 50.5 ± 5.9%; Table 1, Fig. 2a), respectively. Forced-choice tests indicated that the SUITAS also accurately discriminated between high (average of rating 4 and 5) and low (average of rating 1 and 2) anxious arousal in the training and validation datasets (training dataset: accuracy = 100 ± 0%, *P* < 0.001, Cohen’s d = 10.439; validation dataset: accuracy = 100 ± 0%, *P* < 0.001, Cohen’s d = 1.917). Further control experiment and analyses were conducted to rule out the possibility that the results were confounded by color- and motor-related responses potentially involved in the paradigm (see Supplementary Methods, Results and Supplementary Fig. 3–4).

**Fig. 2.**
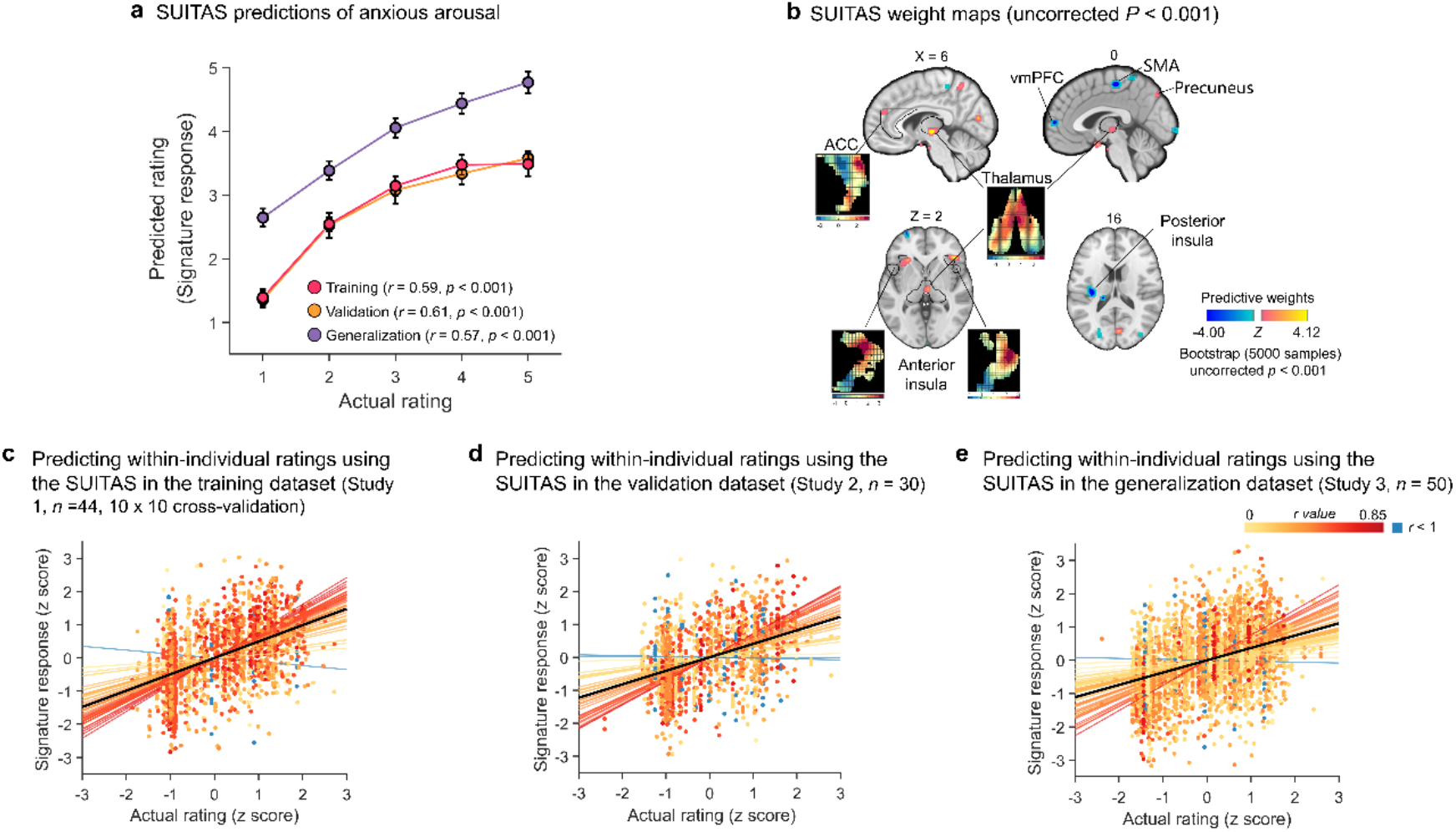
Shock uncertainty-induced threat anticipation signature (SUITAS) model evaluation and weight maps. **a**, Predicted ratings (signature responses) modeled using the anticipatory brain activity (6∼10 s) compared to actual ratings across participants in the training (10 × 10 cross-validated, Study 1), validation (Study 2) and prospective generalization datasets (Study 3). *r* indicates the Pearson correlation coefficient between predicted and actual ratings. Error bars reflect standard error of mean. **b**, The multivariate pattern of fMRI activity predictive of subjective levels of anxious arousal during uncertain threat anticipation (SUITAS weight maps, 10 × 10 cross-validated, Study 1) based on a 5,000 samples bootstrap test. The maps display voxel weights thresholded at *P* < 0.001 uncorrected for display purposes. Inserts show the spatial topography of the unthresholded patterns in the regions previously proposed as core regions of the anxiety network^33^, in particular the ACC, bilateral anterior insula and thalamus. **c**, *Z*-scored actual ratings versus cross-validated predicted ratings (signature response, *z*-scored) within participants in the training dataset (Study 1). Signature response was calculated using the dot product of the SUITAS weight map developed using 10 × 10-fold cross-validation procedure with each single-trial activation map for each participant. **d**, **e**, *Z*-scored actual ratings versus SUITAS predicted ratings (signature response, *z*-scored) within participants in the validation dataset (Study 2) and the prospective generalization dataset (Study 3). Each colored line represents a fitted line for each individual. The black line represents the fitted line across participants. ACC, anterior cingulate cortex; SMA, supplementary motor area; SUITAS, shock uncertainty-induced threat anticipation signature; vmPFC, ventromedial prefrontal cortex.

**Table 1.** Prediction performance (correlation) of the SUITAS on anxious arousal, fear and negative affect ratings.

| Prediction | Anxious arousal | Fear | Negative affect |
| --- | --- | --- | --- |
| Training dataset | 0.59 [0.48, 0.69];<br>0.74 ± 0.01<sup>a</sup> | 0.23 [0.12,0.33];<br>0.47 ± 0.06 | 0.22 [0.15, 0.30];<br>0.49 ± 0.04 |
| Validation dataset | 0.61 [0.50, 0.70];<br>0.77 ± 0.03 | 0.25 [0.05,0.42];<br>0.46 ± 0.08 | 0.26 [0.16, 0.37];<br>0.55 ± 0.06 |
| Generalization dataset | 0.57 [0.48, 0.65];<br>0.75 ± 0.04 | 0.34[0.19,0.47];<br>0.38 ± 0.09 |  |

We applied the SUITAS to subjective anxious arousal, fear and negative affect datasets and calculated the overall (bootstrapped 95% CI) as well as within-individual (mean ± SE) prediction-outcome correlations between the signature responses and the actual ratings. SUITAS, shock uncertainty-induced threat anticipation signature.
<sup>a</sup> indicates cross-validated.

To determine brain regions that reliably contribute to the predictive model and facilitate the interpretation, we performed a bootstrap test through random sampling of participants with replacement from the training dataset with 5,000 iterations. Consistent model weights were thresholded to identify important voxels that reliably contributed to the prediction (uncorrected *P* < 0.001, two-tailed; Fig. 2b for display purposes) which included positive weights in bilateral aINS, thalamus, ACC, superior parietal lobule (SPL), inferior parietal lobule (IPL), dorsolateral prefrontal cortex (dlPFC) and inferior occipital gyrus (IOG), as well as negative weights in ventromedial prefrontal cortex (vmPFC), supplementary motor area (SMA) and posterior insula (pINS).

### Generalization of the SUITAS performance

To test whether the prediction performance of the population-level signature can be generalized to new datasets and paradigms, we applied the SUITAS to a prospective generalization dataset (Study 3, *n* = 50) with different sample and slightly modified UVTA paradigm (e.g., different shock uncertainty baseline, details see Methods) by calculating the SUITAS pattern expressions for each participant (entire anticipation period, one map per rating). The SUITAS could significantly predict anxious arousal ratings and the overall prediction-outcome correlation coefficient was 0.57 (EVS = 19%; bootstrapped 95% CI = [0.48, 0.65], one-tailed permutation test *P* < 0.001; within-participant *r* = 0.75 ± 0.04, mean EVS = 53.4 ± 4.6%; Fig. 2a). Forced-choice test indicated that the SUITAS also accurately discriminated between high and low anxious arousal in Study 3 (accuracy = 94 ± 3%, *P* < 0.001, Cohen’s d = 4.410).

To further determine whether the SUITAS could generalize to other paradigms that encompass an uncertain threat anticipation period, we capitalized on two publicly available datasets acquired during threat conditioning with different MRI systems and scanning parameters in which a visual cue (Study 4, *n* = 59, details see ref.^61^) or auditory cue (Study 5, *n* = 68, details see ref.^62^) (CS+) was paired with a shock on 43% or 33% of the trials, respectively, while a control cue (CS−) was unpaired. We tested whether the SUITAS generalized to distinguish CS+ versus CS−, which is equivalent to uncertain threat versus safe anticipation^63^. Results showed that the SUITAS accurately classified CS+ versus CS− in both datasets (Study 4: accuracy = 79 ± 5%, *P* < 0.001, Cohen’s d = 1.093; Study 5: accuracy = 69 ± 6%, *P* < 0.005, Cohen’s d = 0.490). Overall, the generalizability tests demonstrated that the SUITAS could robustly generalize to new cohorts, paradigms, MRI systems and parameters.

### The SUITAS performance in predicting within-individual anxious arousal

To test whether the population-level SUITAS can track moment-to-moment variations in subjective anxious experience on the individual level, the SUITAS was applied to single-trial activation maps of each participant in Studies 1–3. The SUITAS significantly predicted momentary trial-by-trial ratings within individuals (∼44 trials for each participant in Study 1 and 2, and ∼64 trials for each participant in Study 3, see Methods) with mean prediction-outcome correlation between actual and predicted ratings of *r* = 0.50 (*P* < 0.001, bootstrap test) in Study 1, *r* = 0.41 (*P* < 0.001, bootstrap test) in Study 2, and *r* = 0.37 (*P* < 0.001, bootstrap test) in Study 3 (Fig. 2c, d and e), indicating that the SUITAS was also sensitive to predict within-individual momentary anxious arousal during uncertain threat anticipation.

### A neurofunctional core system for the subjective experience of uncertain threat anticipation

Emotional experience is a highly subjective and individually constructed state with interindividual variations^49^. The mental processes and brain patterns that underlie affective states as well as the construction of the affective state may vary greatly across participants^21^. We further developed within-individual predictive model for each participant using single-trial estimates of brain response (Study 1, ∼44 trials, 10 × 10-fold cross-validated, Supplementary Fig. 5a) and transformed these within-individual patterns to ‘activation patterns’ using structure coefficients (Supplementary Fig. 5b) to determine the core system that reliably and consistently contributed to the prediction and encoded the model (Supplementary Fig. 5c, details see Methods).

The core regions included the aINS, thalamic regions (intralaminar and mediodorsal nuclei), dlPFC, IPL and IOG observed on SUITAS together with anterior midcingulate cortex (aMCC) and parietal regions involved in self-referential processing, e.g., precuneus and posterior parietal cortex (PCC), the putamen and precentral gyrus extending to SMA (FDR *q* < 0.05, retaining positive values, Fig. 3; see also Supplementary Fig. 5c for both positive and negative values). To evaluate spatial distribution of the current results, we mapped the core system onto regions showing activations to uncertain threat versus safe conditions from a recent meta-analysis^33^ (outlined in **Fig. 3**). Visualizing the results indicated a high overlap between the identified networks in the meta-analysis and our core systems (e.g., bilateral aINS, thalamus, dlPFC, SMA) but also emphasized a regional specificity of the core systems, probably reflecting a higher specificity for isolating the subjective experience sub-process. In line with the approach employed in the meta-analysis, we additionally examined the conventional univariate categorical effect (uncertain threat > certain safety anticipation) in our training dataset and observed an extensive and rather process-unspecific bilateral activity spanning the entire insular-cingulate network, lateral frontal and subcortical (e.g., BNST, amygdala, thalamus, and PAG), and occipital regions (FDR *q* < 0.05, see Supplementary Fig. 6).

**Fig. 3.**
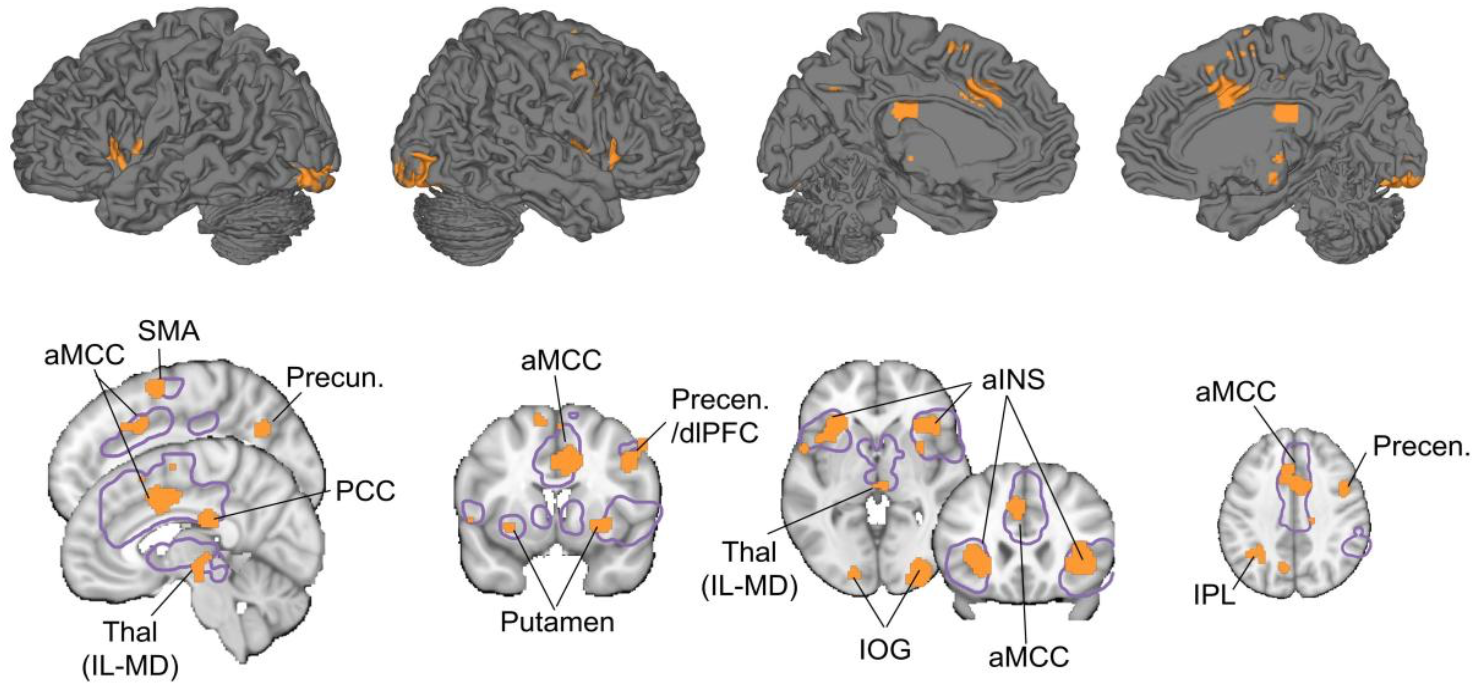
Core brain systems for subjective experience of uncertain threat anticipation. Within-individual core system for subjective experience of threat anticipation under uncertainty using conjunction (retaining positive values) of the within-individual model weight map (one-sample *t* test, FDR *q* < 0.05, based on each participant’s single-trial estimates of brain responses) and the transformed within-individual model encoding map (one-sample *t* test, FDR *q* < 0.05). The violet contour line delineates regions from a meta-analytic activity study comparing uncertain threat versus safe anticipation in healthy individuals^33^. aINS, anterior insula; aMCC, anterior midcingulate cortex; dlPFC, dorsolateral prefrontal cortex; IOG, inferior occipital gyrus; IPL, inferior parietal lobule; PCC, posterior cingulate cortex; Precen, precentral gyrus; Precun, precuneus; SMA, supplementary motor area; Thal (IL-MD), intralaminar and mediodorsal nuclei of thalamus.

### Testing the specificity of the SUITAS against pain, general anticipation and unspecific arousal

The multivariate model might capture processes not specific to anxious arousal but those inherently involved in the UVTA paradigm. We employed a series of analyses with independent datasets (details see Methods and Supplementary Table 1) to determine to which extent the SUITAS captures pain experience, general anticipation and unspecific negative emotional and autonomic arousal. First, we examined whether the SUITAS measured antecedents of the pain response using a publicly available dataset from Wager et al.^53^ (Study 6, *n* = 33). Applying the SUITAS to brain activation maps for thermal pain stimulation periods revealed that the SUITAS distinguished high versus low pain experience (accuracy = 85 ± 3%, *P* < 0.001, Cohen’s d = 0.838, Fig. 4a) less accurately than high versus low anxious experience (accuracy = 94 ± 3%, *P* < 0.001, Cohen’s d = 4.410, Study 3). Second, we asked if the SUITAS could capture the anticipation of non-shock related negative events (loss of money) or the anticipation irrespective of valence (anticipation of a positive event, i.e., gain of money). To this end, we tested the SUITAS on an independent dataset that used a monetary incentive delay (MID) task (https://zib.fudan.edu.cn)^64^ by classifying negative (monetary loss) and positive (monetary gain) versus neutral (no gain or loss) anticipation (Study 7, *n* = 100). The SUITAS could significantly discriminate loss from neutral (accuracy = 65 ± 5%, *P* = 0.004, Cohen’s d = 0.259) but not gain from neutral anticipation (accuracy = 54 ± 5%, *P* = 0.48, Cohen’s d = 0.284, Fig. 4b), although the difference was not significant [(loss vs neutral) vs (gain vs neutral): accuracy = 53 ± 5%, *P* = 0.62, Cohen’s d = 0.021]. These results indicated that the SUITAS might be more effective at identifying negative anticipation than positive anticipation, yet further research is needed to confirm this.

**Fig. 4.**
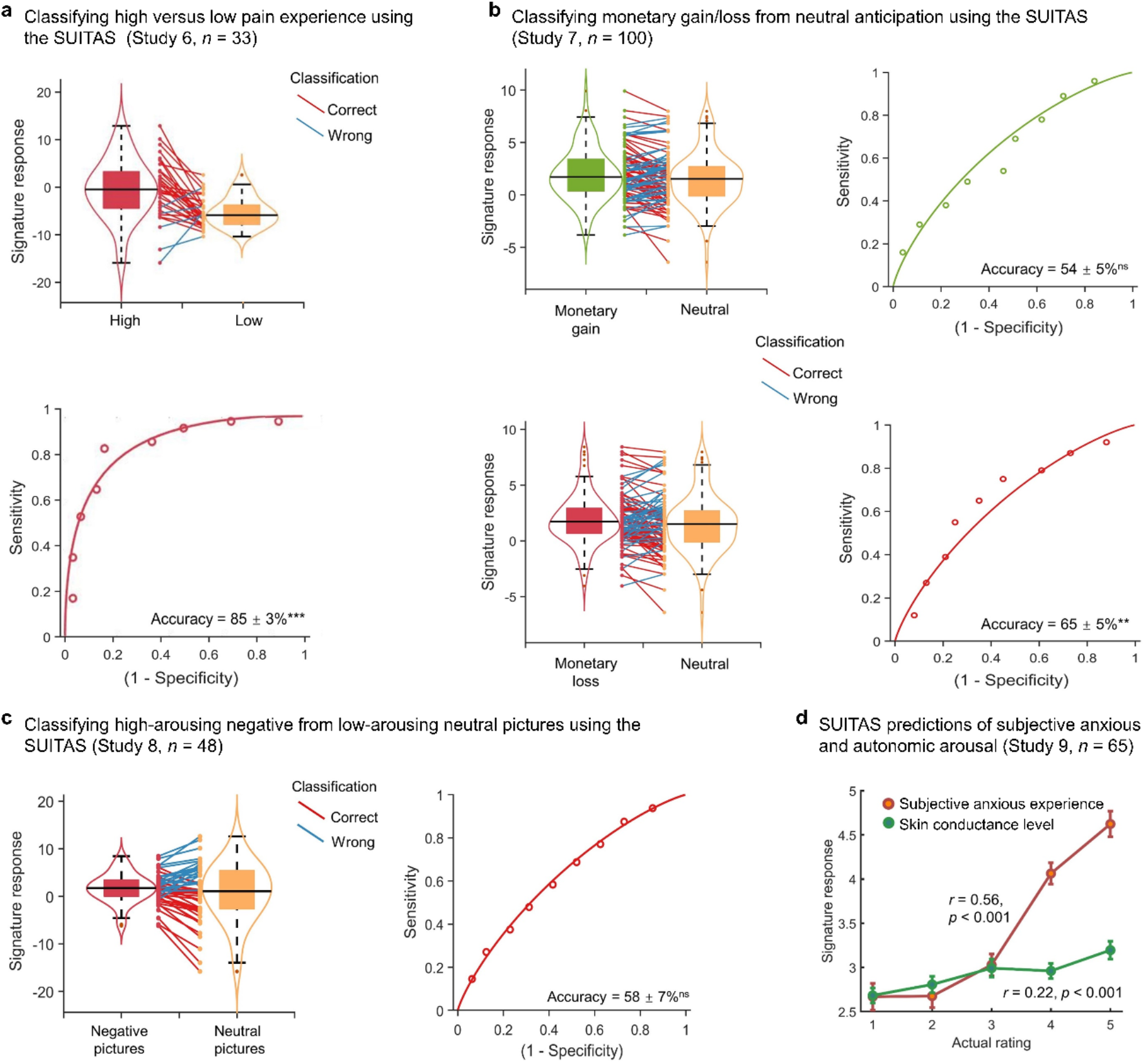
Testing the specificity of the SUITAS against pain, general anticipation and unspecific negative emotional and autonomic arousal. **a,** Classification between high versus low pain stimulation from an independent dataset (Study 6, *n* = 33) using the SUITAS suggests that the SUITAS could predict pain experience yet with lower accuracy and effect size compared to the prediction of subjective anxious experience. The violin and box plots show the distributions of the signature response. The box was bounded by the first and third quartiles, and the whiskers stretched to the greatest and lowest values within median ±1.5 interquartile range. The data points outside of the whiskers were marked as outliers. Each colored line between dots represents each individual participant’s paired data (red line, correct classification; blue line, incorrect classification). **b**, A forced two-choice test of the SUITAS on an independent dataset (Study 7, *n* = 100) using a monetary incentive delay (MID) paradigm reveals that the SUITAS did not distinguish positive (monetary gain) anticipation versus neutral anticipation (top), but could classify non-shock-related negative (monetary loss) versus neutral anticipation (bottom). **c**, Classification between high-arousing negative (disgust) versus low arousing (neutral) pictures from an independent dataset (Study 8, *n* = 48) using the SUITAS suggests that negative arousal did not explain the signature. **d**, Using the SUITAS to predict subjective anxious experience and its physiological arousal (Study 9, *n* = 65) based on subsamples from the validation and prospective generalization datasets. The correlation between actual and predicted ratings (signature response) of subjective anxious experience was more than twice as high as that of skin conductance level. SUITAS, shock uncertainty-induced threat anticipation signature. Performance was shown as accuracy ± SE. \*\*\**P* < 0.001, \*\**P* < 0.01, ^ns^*P* > 0.05.

Third, we accounted for unspecific negative emotional arousal which is inherently associated with uncertain threat anticipation^30,65^ by applying the SUITAS to classify the brain activations during picture viewing period of high-arousing negative (disgust) versus low-arousing neutral visual stimuli (Study 8, *n* = 48). The classification accuracy for high-versus low-arousing stimuli was at chance level (accuracy = 58 ± 7 %, *P* = 0.31, Cohen’s d = 0.235, **Fig. 4c**), suggesting that unspecific negative emotional arousal did not explain the SUITAS. Since arousing pictures may generally rely stronger on visual processing than UVTA, we recomputed the classification accuracy after excluding the occipital lobe of both SUITAS and the test images for arousal. The classification accuracy remained insignificant (accuracy = 54 ± 7%, *P* = 0.67, Cohen’s d = 0.115).

Given that physiological responses are expected to (partly) co-vary with subjective emotional experiences albeit with dissociable neural bases^66^, we finally explored whether the SUITAS predicts subjective anxious experience rather than its physiological correlates by applying the SUITAS to binned brain activation maps for anticipation period of five skin conductance levels (SCL) for each participant. We found that the SUITAS predicted SCL (*r* = 0.216, one-tailed permutation test *P* < 0.001) to a lesser degree than predicting subjective ratings (*r* = 0.556, *P* <0.001; difference in effect size: Δ*r* = 0.34, one-tailed permutation test *P* < 0.001, Fig. 4d, see also Supplementary Results and Supplementary Fig. 7) in the combined validation and prospective generalization datasets (Study 9, *n* = 65), demonstrating that the SUITAS captured autonomic arousal to some extent but with a smaller effect size, which is in line with previous studies on the dissociation between subjective fear experience and its physiological correlates^66^.

Further evidence arguing against the effect of nonspecific negative arousal can be found below (‘Comparing the SUITAS with the predictive models of fear exposure and nonspecific negative affect’). Together with the cross-signature evaluation in the following paragraph, a series of analyses confirm a comparably high specificity of the SUITAS for predicting anxious experience during uncertain threat anticipation.

### Comparing the SUITAS with the predictive models of fear exposure and nonspecific negative affect

We further determined the distinctiveness of our signature from that of fear exposure and general negative affect by comparing the functional and spatial similarities of the SUITAS with the established predictive models of subjective experience of fear (visually induced fear signature, VIFS)^21^ and subjective negative affect (Picture Induced Negative Emotion Signature, PINES)^52^ during picture viewing periods. The prediction performance of the SUITAS on subjective anxious arousal (Study 3: *r* = 0.57, *P* < 0.001) was twice as high as on subjective fear (*r* = 0.23, *P* = 0.015; Δ*r* = 0.34, *P* < 0.001; one-tailed permutation tests) and negative affect (*r* = 0.22, *P* = 0.014; Δ*r* = 0.35, *P* < 0.001; one-tailed permutation tests; Table 1, Fig. 5a) while the VIFS and PINES more accurately predicted subjective ratings of fear and negative affect as compared to the SUITAS (Supplementary Table 2).

**Fig. 5.**
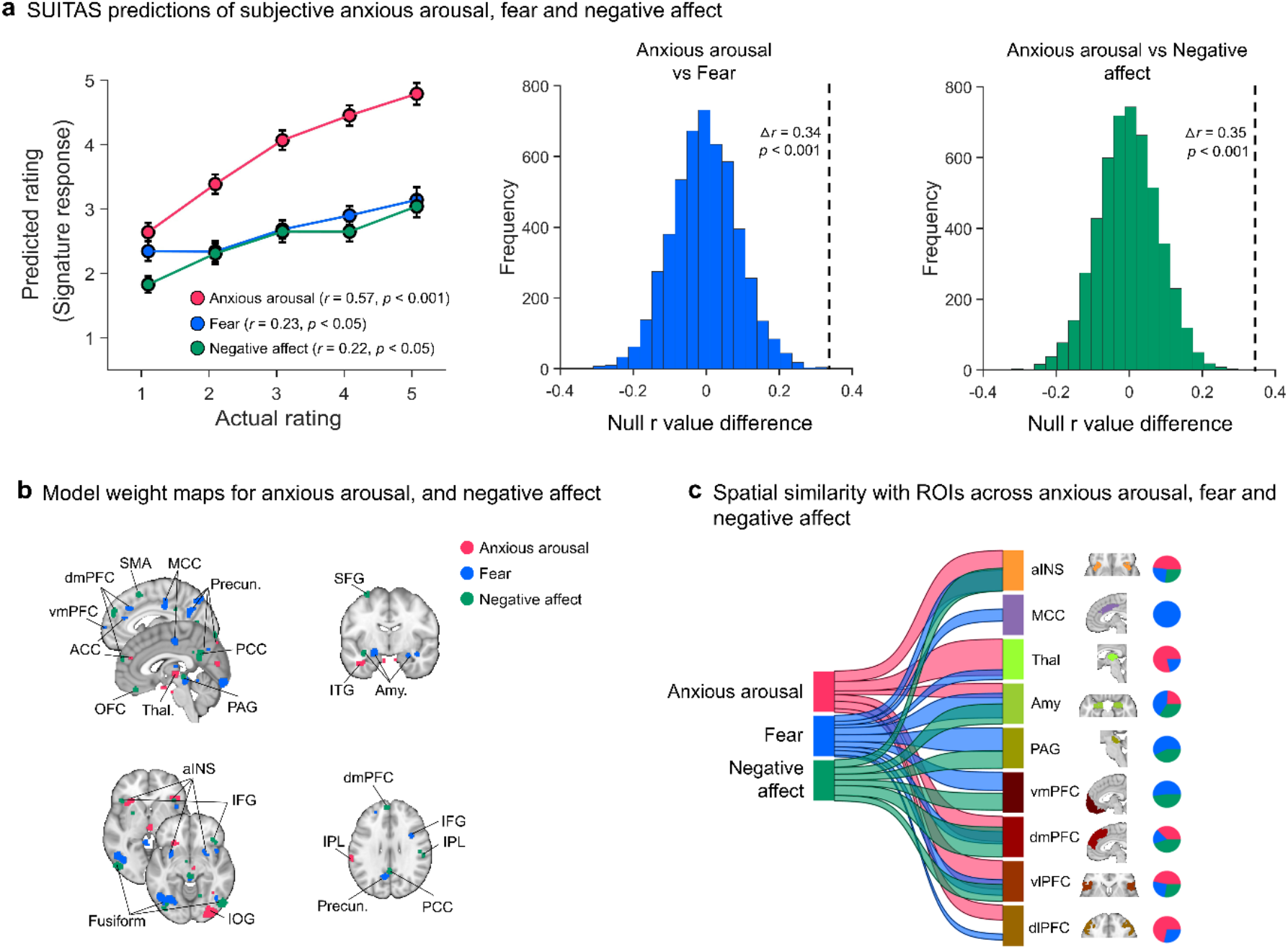
Comparing the SUITAS with the predictive models of fear exposure (VIFS) and negative affect (PINES). **a**, Using the SUITAS to predict subjective experience of fear (*n* = 67, Study 10) and negative affect (*n* = 121, Study 11) based on samples acquired in the previous studies^21,52^. The correlation between actual and predicted ratings (signature response) of anxious arousal (prospective generalization dataset, Study 3) was twice as high as that of fear and negative affect, and the differences of Pearson correlation coefficients (dash lines in the middle and right panel) were significant based on permutation tests of r value difference with 5,000 random shuffles. **b**, Spatial topography of the SUITAS, VIFS and PINES, each thresholded at *P* < 0.001 uncorrected within gray matter and retaining positive values. **c**, River plot depicting the spatial similarity (computed as cosine similarity) between SUITAS, VIFS and PINES weight maps and anatomical labels of predefined ROIs previously linked to negative emotion processing including anxious anticipation, fear exposure and general negative affect. Ribbons are normalized by the max cosine similarity across all ROIs. Maps were thresholded at *P* < 0.001 uncorrected and positive voxels were retained only for similarity calculation and interpretation. Ribbon locations in relation to the boxes are arbitrary. Pie charts show relative contributions of each model to each ROI (that is, percentage of voxels with highest cosine similarity for each predictive map). ACC, anterior cingulate cortex; aINS, anterior insula; Amy, amygdala; dlPFC, dorsolateral prefrontal cortex; dmPFC, dorsomedial prefrontal cortex; IFG, inferior frontal gyrus; IOG, inferior occipital gyrus; IPL, inferior parietal lobule; ITG, inferior temporal gyrus; MCC, midcingulate cortex; OFC, orbitofrontal cortex; PAG, periaqueductal gray; PCC, posterior cingulate cortex; PINES, Picture Induced Negative Emotion Signature; Precun, precuneus; SFG, superior frontal gyrus; SMA, supplementary motor area; SUITAS, shock uncertainty-induced threat anticipation signature; Thal, thalamus; VIFS, visually induced fear signature; vlPFC, ventrolateral prefrontal gyrus; vmPFC, ventromedial prefrontal gyrus.

Applying the SUITAS to classify high versus low levels of subjective anxious arousal, fear and negative affect further confirmed that the SUITAS distinguished high versus low subjective anxious arousal (accuracy = 94 ± 3%, *P* < 0.001, Cohen’s d = 4.410) more accurately than high versus low fear (accuracy = 79.10 ± 5%, *P* < 0.001, Cohen’s d = 0.827) and negative affect (accuracy = 77.14 ± 4%, *P* < 0.001, Cohen’s d = 0.804) in terms of classification accuracy and effect size (see also Supplementary Fig. 8). These results suggest that the SUITAS predicts subjective experience of anxious arousal with high specificity relative to that of fear, subjective negative affect or nonspecific negative arousal. Next, we compared the spatial topography of the SUITAS, VIFS and PINES (**Fig. 5b**). The pattern similarity among the SUITAS, VIFS and PINES weight maps restricted to the gray matter mask suggests that these models exhibited weak positive spatial correlations on the whole-brain level (SUITAS versus VIFS: *r* = 0.07; SUITAS versus PINES: *r* = 0.05; VIFS versus PINES: *r* = 0.08, one-tailed permutation test all *P* < 0.001). The Pearson correlations decreased after thresholding the model weights at both uncorrected *P* < 0.01 and uncorrected *P* < 0.001 (see Supplementary Results for details). To test to what extent the similarity or distinction of performance depended on the contribution of the visual cortex, we retrained these models excluding the occipital lobe and then compared the functional and spatial similarities and the results remained consistent (see Supplementary Results and Supplementary Fig. 9), suggesting little contribution of visual cortex in the differentiation among predictive models of threat anticipation, fear exposure and negative affect.

We further used a river plot to illustrate the spatial similarity between predictive weights of these models (uncorrected *P* < 0.001) and a set of *a priori* regions of interest (ROIs, see Supplementary Table 3) previously linked to negative emotion processing including anxious anticipation, fear exposure and general negative affect^33,67,68^. As shown in Fig. 5c, each ROI encoded at least one model with different contributions: thalamus, dlPFC, aINS and vlPFC were contributed most by SUITAS, whereas MCC, amygdala, PAG and vmPFC were contributed most by the VIFS and dmPFC was contributed most by the PINES. Summarizing, the double dissociation of predictive performance and the distinct spatial topography suggest that these signatures show distinct representations in predicting their corresponding subjective emotional states.

### Single subsystems are not sufficient to predict subjective experience of threat anticipation under uncertainty

Previous ‘structure-centric’ theories proposed that emotions are localizable in single brain regions or networks^18,69–71^. To determine whether subjective emotional experience of threat anticipation under uncertainty can be reduced to specific systems, we re-trained the predictive models in (1) five *a prior* regions considered as ‘classical’ anxiety systems ^13,33^; (2) salience network hubs associated with anxiety and anxiety disorders^72^; (3) cortical network related to conscious emotional experience^18^; (4) seven large-scale functional networks^73^, and tested them on training, validation and generalization datasets. Isolated regions (i.e., aINS, ACC, thalamus and BNST, but not PAG) and networks could predict subjective ratings, however, with considerably smaller effect size as compared to the whole-brain model (see Supplementary Results and Supplementary Fig. 10a). To control for the potential effect of the numbers of features in the prediction analyses (i.e., whole-brain model uses much more features/voxels), we randomly selected certain numbers of voxels from a uniform distribution spanning the whole brain or aforementioned networks. The asymptotic predictions with sampling from all brain systems were consistently higher than sampling the same number of voxels from the individual networks when exceeding 1,000 voxels (Supplementary Fig. 10b), suggesting that the whole-brain model has much larger effect size than those using the same number of features from single networks.

In line with accumulating evidence^21,52^, the above results underscore that subjective experiences of uncertain threat anticipation require a distributed neurofunctional representation.

## Discussion

Aberrant anticipatory responding of uncertain threat is central to anxiety-related disorders^2,11^. We here developed an uncertainty-variation threat anticipation paradigm and utilized multivariate predictive modeling to systematically determine an accurate, robust, generalizable and specific fMRI-based neuro-affective signature for the subjective experience of threat anticipation under uncertainty using several independent datasets in healthy adults. The developed SUITAS was predictive of the level of uncertainty-induced subjective anxious experience on both the population and individual level. The signature showed a robust generalization across cohorts and paradigms and was less or not sensitive to processes inherently interwoven with the paradigm, including pain, positive anticipation (but negative anticipation), or unspecific negative emotional and autonomic arousal, suggesting a comparably high specificity for predicting anxious anticipation of aversive events. Comparison with established decoders for subjective fear and negative affect indicated a certain extent of cross-prediction, suggesting that the decoder may partly capture a common underlying process, e.g. conscious emotional experiences or interpretation^74^, but the considerably higher effects size for the target emotional state also suggests distinguishable neural representations of threat anticipation, fear exposure and negative affect. Consistent model weights associated with subjective anxious arousal were observed in the aINS, thalamus, ACC, somatosensory cortices and IOG, while the within-individual models additionally revealed consistent contributions of aMCC, precuneus, PCC, putamen, precentral gyrus and SMA to predict momentary trial-by-trial variations. No single brain region or network was sufficient for accurately predicting anxious experience, underscoring that conscious emotional experiences require a distributed neural representation^10,21,33,52,75,76^. The neuro-affective signature provides a promising neuroimaging biomarker for subjective anxious experience during uncertain threat anticipation which can facilitate rapid and accurate evaluation of novel interventions on anxiety (see e.g. ref.^77^) with potential for clinical translation.

Uncertainty about potential threat has long been considered as key candidate mechanism underlying anxiety^2,78,79^. However, how varying levels of uncertainty impact the subjective experience of threat anticipation remained unknown. Our UVTA paradigm based on the established TOS paradigm - but manipulating different aspects of anticipatory uncertainty - successfully induced sufficient and varying intensity levels of anxious anticipation. Moreover, individuals high in intolerance of uncertainty reported higher anxious arousal in the uncertain conditions of the UVTA task. These results together demonstrated uncertainty as candidate mechanism underlying the experience of anxious arousal and allowed us to determine the underlying neuro-affective signature.

Employing a pattern recognition-based machine learning technique that has been previously successfully employed to other subjective emotional experience domains^21,52,53,74,80,81^ allowed us for the first time to identify a robust and process-specific predictive pattern for the subjective experience of uncertain threat anticipation and segregate it from other emotional states. Importantly, out-of-sample predictions in two independent samples (study 2 and 3) protected against overfitting and showed the robustness and generalizability of the brain-wide model^82^. The generally higher signature response across rating levels of the prospective generalization dataset might attribute to the higher baseline of uncertainty related to including a higher proportion of uncertain trials of the UVTA task in Study 3 (see Fig. 2a).

The SUITAS provides evidence that the subjective experience of uncertain threat anticipation involves distributed neural systems^75^, which resembles current observations in animal models that the response to uncertain threats recruits a distributed array of interconnected neural ensembles^10^. Further analyses demonstrated that no single region or network was sufficient for predicting anxious experience, which aligns with predictive modeling results for other subjective emotional experiences^21,52^ and argues against the traditional structure-centric view^70,71^ but rather suggests a constructionist theory of emotion^48,49^. The core system is partly consistent with recent meta-analyses on induced anxiety-associated brain activity^33^ and structural alterations in anxiety-related disorders^83^. The aINS, thalamus and aMCC are key regions of the cingulo-opercular network^84,85^, which partly resembles the salience network^86^. These regions in concert not only support the detection of threatening signals^85^ but also interoceptive processes, emotional awareness of negative affect, and autonomic activity^87–91^. The dlPFC, IPL and precuneus constitute core nodes of the fronto-parietal network which exerts top-down feedback control and is involved in constructing conscious experiences and self-related processes^18,92^. These regions together promoted the subjective experience of anxious arousal on both the population and individual level, which may suggest that conscious experience of uncertain threat anticipation is a constructed state that encompasses different functional modules^48,49^.

The aINS has long been suggested to represent subjective feelings from the body and emotional awareness across emotional domains including anger, fear, sadness, happiness, disgust and aversion^87^. Moreover, the aINS is involved in uncertainty and risk processing besides its established role in perception of interoceptive and subjective feeling states^2,93^. For example, the aINS plays a critical role in the anticipation of uncertain threats relative to certain threats^45,94,95^. Bilateral aINS activity has also been associated with risk prediction in reward anticipation under uncertainty^96^. The aINS exhibits reciprocal connections with the aMCC^97,98^, a region proposed to integrate information about punishment, interoceptive and subjective emotional states to exert control in the face of uncertainty^90^. Our results therefore support the critical roles of aINS and aMCC in detecting, interpreting and reacting to salient internal and environmental changes in the face of uncertainty which is central to anxiety.

The thalamus consists of different nuclei that serve various functions including relaying sensory and motor signals, memory, attention as well as the regulation of consciousness, alertness and emotion^99,100^. The present study robustly identified intralaminar and mediodorsal thalamic nuclei as critical modules predictive of subjective anxious arousal on both the population and individual level, which may be related to their function in the integration of information across multiple cortical circuits that influences the conscious experience of anxious arousal during uncertain anticipation of the future^101^.

Importantly, although the aINS, thalamus and aMCC are key nodes of the salience network^85,86^, our control analyses suggested that the SUITAS did not respond to salience or unspecific negative emotional and autonomic arousal alone (Study 8 and 9). Moreover, none of these regions or other networks was sufficiently predictive of subjective anxious arousal, suggesting that the subjective experience during uncertain threat anticipation requires a distributed and brain-wide engagement to support the idiosyncratic and complex emotional experience. Notably, although the BNST has been suggested to play a prominent role in animal and human models of anxiety^14,30,102^, we did not find BNST in the signature predictive of anxious anticipation which may suggest that BNST representation did not code for subjective experience despite its role in automatic processing of uncertain threats^30–32^.

An important contribution of the current study is the comparison of the neural representation of subjective experience of uncertain threat anticipation with that of subjective fear (Study 10) and negative affect (Study 11). The conceptual and neural differentiation of fear (acute threat), anxiety (potential threat) and nonspecific negative affect has long been debated in neurobiological models of emotion^17,21,30,33,83,103^, animal models^11,15,104,105^ and neuropsychiatric models such as the RDoC framework^6,7^. Rodent models suggest that defensive and physiological responses related to fear or anxiety are mediated by distinct neural substrates, i.e., central nucleus of the amygdala and BNST, respectively^11^. In contrast, human fMRI findings on functional dissociations between fear responses and uncertain threat anticipation have been inconsistent^29,30,33,36^. Recent evidence from meta-analysis of human fMRI studies suggests that ‘fear’ (conditioned versus unconditioned stimulus) and ‘anxiety’ (uncertain threat versus safe anticipation) may engage a partly overlapping circuit including BNST, MCC, aINS and PAG^33,106^. However, the mass univariate and categorical contrast approach does not permit to clearly segregate the neural systems underlying different mental processes such that common neural circuits may alternatively reflect hard-wired defensive behaviors or physiological responses as well as the generally increased arousal that characterize both fear and anxiety. The present study demonstrates fine-grained distributed neural patterns can, to a certain extent, segregate subjective experiences of uncertain threat anticipation, fear exposure and negative affect functionally and spatially such that the classic ‘emotional’ regions showed different levels of involvement in representing different emotion domains (see Fig. 5c), which suggests that these regions may to a certain extent encode anxious arousal, fear and negative emotions in distinguishable neural representations.

Several important caveats should be taken into account when interpreting the results of the present study. First, the developed model was based on uncertain anticipation of electric shocks, and the generalization datasets also used shocks as aversive stimuli. Including datasets with other types of uncertain threats would allow for a more robust test of the generalizability of our brain model. Second, the parameter settings (C and epsilon) of the SVR were based on previous studies with the similar purpose. However, a grid search is invited to determine the optimal combination of hyperparameters that allows to yield the highest performance in future studies. Moreover, there might be differences in the task design (e.g., with rating or not) and experimental parameters (e.g., stimulus type, durations) between our original datasets and the external datasets for testing the specificity of the SUITAS. Although we employed control experiments and analyses to validate that the results were not explained by these potential confounding factors, datasets with comparable task design and stimulus type are needed in future studies for the specificity test. Lastly, the sample size for the training dataset was based on previous studies^80^. While researchers recently proposed that testing generalizability of model in novel samples could provide evidence for a robust neuromarker without large sample size^107^, research on a priori power and sample size estimations for multivariate neuroimaging models in task-fMRI will critically advance the field.

Together, the current study provided a comprehensive brain-level description of subjective experience of threat anticipation under uncertainty. The sensitive, generalizable and specific signature has potential to be tested prospectively in future studies with different experimental settings and populations (e.g., patients) and promote clinical translation.

## Methods

### Participants

A total of 124 healthy young participants (64 females, age range: 18– 33) participated Studies 1–3 to develop (Study 1, *n* = 44), validate (Study 2, *n* = 30) and prospectively test the generalizability (Study 3, *n* = 50) of a multivoxel-pattern based predictive model of subjective experience of uncertain threat anticipation. All studies were approved by the local ethics committee at the University of Electronic Science and Technology of China and were in accordance with the latest revision of the Declaration of Helsinki. All participants provided written informed consent and were remunerated for their participation. Data from Study 1 and Study 2 were acquired in June 2021 (T1) using the same experimental design and scanning parameters at the same study site, while Study 3 was conducted 10 months later (April 2022, T2) with a slightly different experimental design (e.g., more experimental trials, different threat probability and uncertainty baseline, details see Paradigm and procedures) and was a part of our ongoing project examining cue-based anticipation of multimodal affective input. We randomly selected 60% of the participants from T1 as the training dataset to develop the brain model (Study 1: *n* = 44, 23 females, mean ± SD age = 22.07 ± 2.50 years). The remaining participants comprised the validation dataset (Study 2: *n* = 30, 14 females, mean ± SD age = 22.47 ± 2.76 years). Data from T2 were used as the prospective generalization dataset (Study 3: *n* = 50, 27 females, mean ± SD age = 20.08 ± 2.22 years) to test if the brain model identified in Study 1 can be prospectively applied to independent data and generalized to different shock probability and uncertainty environments. Participants were excluded if they reported a current or history of neurological, psychiatric or major physical disease, psychotropic medication, substance abuse, MRI contraindications and any prior participation in experiments with electric stimulation.

### Paradigm and procedures

*Study 1 and Study 2.* In Study 1 and Study 2, participants completed an uncertainty-variation threat anticipation (UVTA) task while undergoing fMRI acquisition. Before the start of the actual experiment, participants were instructed that they would anticipate electric shocks with varying levels of uncertainty and provide ratings of the anxious levels they experienced during the anticipation period at the end of each trial. Prior to MRI data acquisition, a shock calibration was implemented during which a highly aversive but not unbearable shock level was determined for each participant. Participants rated subjective shock discomfort on a scale from 1 (not at all painful) to 5 (painful and difficult to tolerate) to reach a level of 4 (painful but not unbearable). The shock calibration procedure was repeated in the middle of the four experimental runs to avoid habituation. 75% of participants in Study 1 and 70% of participants in Study 2 adjusted the level of electric stimulation during the recalibration. Electric shocks were delivered to the underside of the left wrist using a Biopac STM100C (Biopac Systems Inc., Goleta, CA). The UVTA paradigm used an event-related design consisting of four conditions repeatedly presented over four runs in a pseudorandom order with no more than two consecutive trials of the same condition. The four conditions corresponded to four different uncertainty levels from certain safety, low, medium to high threat uncertainty modulated by different combinations of event (shock), temporal (duration) and number of shock uncertainty. Low, medium and high uncertainty conditions were cued by a colored lightning bolt (blue, purple and red, respectively) in the center of the screen and a border of the same color whereas the certain safety condition was indicated by a white isosceles triangle and a white border (Fig. 1a). Prior to the task, participants were told about the contingency between cues and outcomes, e.g. that the blue cue would last 8 s and two consecutive shocks might occur immediately after the cue presentation (low threat uncertainty: event-only uncertainty); the purple cue would disappear any time between 0 to 16 s and two consecutive shocks might occur immediately after the cue presentation (medium threat uncertainty: event and temporal uncertainty); the red cue would disappear any time between 0 to 16 s and two consecutive shocks might occur or three shocks might occur either in a row or separated into two consecutive shocks immediately after the cue presentation and one shock after the subsequent fixation epoch (high threat uncertainty: event, temporal and number uncertainty); no shocks would be administered after the white cue lasting between 0 to 16 s (certain safety). The exact shock probability for low, medium and high uncertainty conditions was not stated to the participants and they were only told that the likelihood of shock outcomes of these three conditions was equivalent. The shock probability was predetermined to 60% for each uncertain threat condition. The purple, red and white cues in actuality ranged between 6 and 10 s (avg 8 s) and both medium and high uncertainty conditions contained ‘fast’ trials with 3 s of cue presentation and ‘slow’ trials with 13 s of cue presentation pseudo-randomly to increase the credibility of the instruction that the cues could disappear any time within 16 s after which shocks might occur. Each run contained 5 valid trials per condition and 4 dummy trials (2 fast trials and 2 slow trials, shock probability: 50% throughout all runs), resulting in 24 trials in total in each run. The dummy trials were not included in the behavioral and fMRI analysis. Participants were asked to retrospectively rate their subjective level of anxious arousal during the anticipation period (6∼10 s) from 1 (no anxiety) to 5 (extreme anxiety). Stimuli were presented via E-Prime 2.0 (Psychology Software Tools, Sharpsburg, PA). Note that the varying levels of uncertainty conditions were deployed to induce different levels of subjective anxious feelings and the behavioral rating patterns confirmed that our paradigm successfully evoked varying levels of anxiety between conditions (Fig. 1b and Supplementary Fig. 1).

MRI data were collected using a GE Discovery MR750 3.0T system (General Electric Medical System, Milwaukee, WI, USA). Functional images were acquired with an interleaved T2*-weighted gradient echo-planar imaging sequences (40 slices; repetition time (TR) = 2000 ms; echo time (TE) = 30 ms; slice thickness = 3.8 mm; spacing = 0.6 mm; field of view (FOV) = 200 × 200 mm; flip angle = 90°; matrix size = 64 × 64; voxel size = 3.125 × 3.125 × 3.8mm). High-resolution whole-brain T1-weighted images were additionally acquired to improve spatial normalization (3D spoiled gradient echo pulse sequence; 154 slices; TR = 6 ms; TE = 3 ms; slice thickness = 1mm, FOV = 256 × 256 mm, acquisition matrix = 256 × 256, flip angle = 8°, voxel size = 1 × 1 × 1 mm).

*Study 3.* Study 3 implemented a modified UVTA task. The differences were that the total number of trials and the proportion of uncertain threat trials increased (each run contained 29 trials, 8 trials per low, medium and high uncertainty conditions, 4 trials per certain safety condition and 1 ‘fast’ trial), which resulted in increased statistical power and a different shock uncertainty baseline. The shock probability was predetermined to 50% for each uncertain threat condition and the ‘fast’ trial for each run belonged to a different condition and was always presented in the first trial with shocks. The shock calibration procedure was the same and 70% of the participants adjusted the level of electric stimulation in the middle of the experimental runs. A chi-square test confirmed no habituation effects of shock pain on anxious experience over the course of the paradigm (*χ*^2^(3) = 16.21, *P* = 0.18; Study 1–3). Participants were asked to retrospectively rate their subjective level of anxious arousal during the anticipation period (6∼10 s) from 1 (not anxious at all) to 5 (extremely anxious). Imaging data acquisition were identical to Study 1 and Study 2 except that the participants in Study 3 moreover performed additional emotional picture tasks after UVTA.

### fMRI preprocessing and analysis

The fMRI data were preprocessed and analyzed using Statistical Parametric Mapping (SPM12, https://www.fil.ion.ucl.ac.uk/spm/software/spm12/) and the procedures for preprocessing and general linear model (GLM) analyses were the same for Study 1, 2 and 3. The first five volumes of each run were discarded to allow for magnetic field equilibration. Prior to preprocessing, image intensity outliers were identified using CanlabCore tools (https://github.com/canlab/CanlabCore, for details, see ref.^21^). Each time-point identified as outliers was included in the first-level model as a separate nuisance covariate. The remaining functional images were correct for slice timing differences, head movements, co-registered with the T1-weighted structural images, normalized to Montreal Neurological Institute (MNI) standard template (interpolated to 2 × 2 × 2 mm voxel size) and spatially smoothed with an 8-mm full-width at half maximum Gaussian kernel (For details, see ref.^21^).

Preprocessed images were subjected to a first-level GLM using SPM12 for prediction analysis. The four runs were temporally concatenated beforehand to ensure that there were enough trials for each rating. The model included five separate boxcar regressors time-locked to the anticipation period (6∼10 s) corresponding to each rating (i.e., 1–5), which allowed us to model brain activity in response to each anxious level separately. Note that we only included the anticipation periods after which no shocks occurred to obviate any potential interference of the shock pain^108–110^. Accordingly, the anticipation periods followed by shocks, the ‘fast’ and ‘slow’ trials as well as missed trials, were treated as regressors of no interest. The outcome period was also modeled with a boxcar regressor to examine the effects of shock delivery. To model any effects related to motor activity, one boxcar regressor indicating the rating period was included. The two fixation-cross epochs served as implicit baseline. All regressors of interest were convolved with a double gamma canonical hemodynamic response function (HRF). Additionally, 24 head motion parameters (6 realignment parameters demeaned, their derivatives and the squares of these 12 regressors), indicator vectors specifying ‘spikes’ by framewise displacement (FD) that had deviations larger than 0.50 mm^111^ and outlier time points identified from image intensity (see above for details) were treated as nuisance regressors.

### Developing the brain model

Using the training dataset (Study 1, *n* = 44), we developed a whole brain neural signature (gray matter masked) predictive of subjective anxious experience (i.e., SUITAS) by training a support vector regression (SVR) model. We first concatenated each participant’s data across runs and then used individual first-level GLM images (one per rating of no-shock trials for each participant) across participants as features to predict participants’ ratings of the anticipation periods (Fig. 1c). The SVR was performed using a linear kernel function (*C*=1) with epsilon setting to 0.1 in the Spider toolbox (http://people.kyb.tuebingen.mpg.de/spider) in line with previous studies^20^.

### Model evaluation

To evaluate the model performance and minimize overfitting, we used 10 × 10-fold cross-validation within the training dataset in Study 1 and applied the brain model (i.e., SUITAS) to new individuals in Study 2 (Fig. 1c). The cross-validation procedures were repeated ten times by producing different splits (10 subsamples) in each repetition. Beta images from 9 subsamples were trained to predict the anxiety ratings, and then the performance was tested on the holdout participants (1 subsample). To obtain unbiased estimates of the model performance, we predicted subjective ratings in new individuals from the validation dataset (Study 2, *n* = 30) by calculating the signature responses using a dot product of the SUITAS weight map with each vectorized activation map (one per rating). To provide an interpretable effect size metric, we calculated Pearson correlations between the predicted ratings and the actual ratings across participants for both Study 1 and Study 2, and the statistical inference was determined using a permutation test with 5,000 random shuffles only for Study 2 given that cross-validated permutation test is very time consuming. Explained variance score (EVS) was computed to indicate the overall prediction error (for similar approaches see ref.^21^). We additionally assessed the classification accuracy between high (average of rating 4 and 5) versus low (average of rating 1 and 2) levels of subjective anxious ratings using two-tailed forced-choice tests from receiver operating characteristic curves.

### Generalization of the SUITAS

We first tested whether SUITAS could predict subjective anxious ratings in new individuals from Study 3 (*n* = 50) who were not included in the initial model training and validation and underwent a modified UVTA (Fig. 1d). The independent dataset in Study 3 could provide a good test for the model generalizability and also allow us to obtain unbiased estimates of the model sensitivity. Similarly, pattern responses were estimated for each test participant by computing the dot product of SUITAS pattern with the participant’s vectorized activation map. The calculations for Pearson correlation, permutation test, EVS and classification accuracy were the same as Study 2. Moreover, we used two publicly available datasets (Study 4, *n* = 59 from our previous study using a visual threat-conditioning paradigm^61^; Study 5, *n* = 68 from a previous study using an auditory threat-conditioning paradigm^62^; details see Supplementary Methods and Supplementary Table 1) to further test the generalizability of the SUITAS in distinguishing uncertain threat versus safe anticipation across cohorts, paradigms, MRI systems and scanning parameters.

### Within-individual prediction

To further determine the sensitivity of the population-level model (i.e., SUITAS) in tracking within-individual variation in anxious ratings, we estimated single-trial responses using a GLM design matrix with separate regressors for each trial (no shock) in Study 1, 2 and 3, resulting in ∼44 trials for each participant in the training and validation datasets and ∼64 trials for each participant in the prospective generalization dataset. The SUITAS was next applied to the single-trial activation maps to obtain the signature responses, which were finally correlated with the true ratings for each participant separately. Note that we used 10 × 10-fold cross-validation to obtain less biased test results for each participant in Study 1, whereas the within-individual prediction in Study 2 and Study 3 did not use cross-validation procedure because they were not included in the training of the model. The correlation coefficients were first *r* to z transformed and then were averaged to obtain one mean correlation value. Bootstrap tests (5,000 iterations) were used to test whether the distributions of within-individual prediction–outcome correlations were significantly higher than zero.

### Identifying a core system consistently involved in subjective experience of anxiety

To identify the important brain regions for subjective experience of uncertain threat anticipation, we located the consistent voxels across participants that (1) reliably contributed to the model prediction (i.e., model weights) and (2) were associated with subjective anxious arousal (i.e., model encoding). Previous studies have suggested that both model weight (‘betas’) and model encoding maps (‘structure coefficients’) are necessary to interpret the model^74,112^. We therefore obtained both model weight maps and model encoding maps at the individual level and then calculated their conjunction (i.e., significant voxels in both maps) to determine the core brain system for subjective experience of threat anticipation under uncertainty.

To do this, we first ran a separate prediction analysis (linear SVR with C= 1) for each participant in the training dataset (Study 1, *n* = 44) using their single-trial beta images (∼44 trials, 10 × 10-fold cross-validated). A one-sample *t* test (FDR *q* < 0.05) across within-individual patterns was performed to evaluate the consistency of each weight for every voxel in the brain. Next, model encoding (‘structure coefficient’) maps were computed for each participant by transforming the within-individual predictive pattern using the following formula: *A* = *cov*(*X*) × *W* × *cov*(*W*^T^ × *X*)^-1^, where *A* is the reconstructed activation pattern, *cov*(*X*) is the covariance matrix of individual data, *W* is the SUITAS predictive weights, and *cov*(*W*^T^ × *X*) is the covariance matrix of the latent factors. Structure coefficients map individual voxels to the overall multivariate model prediction from backward model to forward model and the significant brain regions were determined by one-sample *t* test (FDR *q* < 0.05) on the within-individual encoding maps where the voxel-wise activity correlates with the model prediction. The conjunction (FDR *q* < 0.05, preserving positive values) of the thresholded second-level within-individual model weight map (one-sample *t*-test) and model encoding map (one-sample *t*-test) was considered as the core system map. The regions were moreover mapped onto the results from a recent meta-analysis on uncertainty induced threat anticipation that included categorical comparisons of fMRI activation between unpredictable threat and safe anticipation in healthy individuals^33^.

### Testing the specificity of the SUITAS against pain, anticipation and arousal

To examined whether the SUITAS specifically captures anxious feelings, we evaluated the extent to which the SUITAS was sensitive to pain experience, general positive/negative anticipation and unspecific negative emotional arousal involved in the paradigm by testing the SUITAS on four independent datasets (Study 6, *n* = 33 from previous studies using a thermal pain paradigm^53,113^; available at https://figshare.com/articles/dataset/bmrk3_6levels_pain_dataset_mat/6933119; Study 7, *n* = 100 from an ongoing study using a monetary gain/loss anticipation paradigm - MID task^114–^ to measure reward processing, https://zib.fudan.edu.cn; Study 8, *n* = 48 from an ongoing study using an emotional picture paradigm and was a subsample from Study 3, Fig. 1d; Study 9, *n* = 65 from Study 2 and 3 in which the skin conductance data was recorded; see also Supplementary Methods and Supplementary Table 1 for details), respectively. For Study 6, we used the SUITAS to classify the pain stimulation periods corresponding to high (5 and 6) versus low (1 and 2) pain ratings after each trial. For Study 7, we applied the SUITAS to distinguish anticipation of monetary gain versus neutral condition and monetary loss versus neutral condition where signature responses were compared for two conditions. For Study 8, we applied the SUITAS to classify exposure towards pre-selected high-arousing disgust versus low-arousing neutral pictures that were presented in an event-related design during fMRI acquisition (details see Supplementary Methods). Classification accuracy was calculated from receiver operating characteristic curves using forced-choice classification and *P* values were calculated using two-tailed independent binomial tests for Study 6–8. Moreover, we tested whether the SUITAS could specifically predict subjective anxious feelings rather than its concomitant skin conductance level using subsamples of Study 2 (*n* = 22) and Study 3 (*n* = 43) who had complete skin conductance data (Study 9, details see Supplementary Methods and Supplementary Table 1).

### Comparing the SUITAS with predictive models of subjective fear and negative affect

To test whether or to which extent The SUITAS was distinct from neural representations of fear exposure and negative affect, we compared the functional and spatial similarity of the SUITAS with the subjective fear signature VIFS from our previous study (Study 10, *n* = 67; details see ref.^21^) and the subjective negative affect signature PINES from Chang et al. (Study 11, *n* = 121; details see ref.^52^) (Fig. 1d, see also Supplementary Methods for details). Specifically, we (1) applied the SUITAS to the datasets of VIFS and PINES, and vice versa (i.e., applied the VIFS or PINES to the datasets of SUITAS) to examine the prediction performance, and (2) compared the spatial topography of the SUITAS, VIFS and PINES patterns by calculating the spatial correlation among these predicted weights (unthresholded and thresholded at uncorrected *P* < 0.001) and computed cosine similarity between these signatures and predefined ROIs documented in previous studies as regions showing preferential activation to anxiety^33^, fear^67^ and negative affect^68^. To exclude the visual processing effect, we retrained these models excluding the occipital lobe and compared the performance of SUITAS with VIFS and PINES (see Supplementary Results and Supplementary Fig. 9).

### Models using features from local regions or networks

To examine whether the subjective anxious experience was reducible to activation patterns in any single brain region or network, we re-trained SVR models (10 × 10-fold cross-validated) restricted to local brain regions (e.g., ACC, aINS, thalamus, BNST, etc.) and networks (e.g., seven large-scale resting-state function networks^73^, salience network^72^ and consciousness network^18^) which have been previously suggested to be associated with anxious anticipation. Anatomical parcellations and the corresponding atlases used to select ROIs are detailed in Supplementary Table 3.

## Acknowledgements

This work was supported by the National Natural Science Foundation of China (NSFC, Grant No. 32250610208; 82271583), the Ministry of Science and Technology (MOST2030, Grant No. 2022ZD0208500) and the National Key Research and Development Program of China (Grant No. 2018YFA0701400). The funders had no role in study design, data collection and analysis, decision to publish or preparation of the manuscript. Any opinions, findings, conclusions or recommendations expressed in this publication do not reflect the views of the Government of the Hong Kong Special Administrative Region or the Innovation and Technology Commission.

## Author contributions

X.Q.L and B.B. conceived and designed the study. X.Q.L and G.J.J. collected the data. X.Q.L analyzed the data. X.Q.L and B.B. drafted the manuscript. F.Z., K.M.K, D.Y., S.X., T.J., X.Z., J.Z., J.F. contributed data and important intellectual input on the manuscript.

## Competing interests

The authors declare no competing interests.

## Additional information

The online version contains supplementary material.

## Notes

### Competing Interest Statement

The authors have declared no competing interest.

