## Supplements for "A neural signature for the subjective experience of threat anticipation under uncertainty"

**This Word file includes:**

1. Supplementary Methods

2. Supplementary Results

3. Supplementary Figures 1-10

4. Supplementary Tables 1-3

**Supplementary Methods**

Control experiment: number rating anticipation

To rule out the possibility that the developed model SUITAS was confounded by the color of the cue or the motor-related response, we designed and conducted an fMRI experiment aimed at controlling for the effect of color or preparatory motor response. Participants were shown five different colored numbers from 1-5, each with a specific color (Supplementary Fig. 3). They were required to anticipate the number that they were going to rate during the number presentation stage and rate according to what they previously saw on a scale ranging from 1 to 5 at the end of each trial. The duration for each phase was kept the same as the UVTA paradigm. Stimuli were presented using the E-Prime software (Psychology Software Tools, Sharpsburg, PA).

We collected an independent dataset (n = 20, 8 females, 18-26 years, 21.30 ± 2.74) to test potential confounding effects of the processes on the SUITAS. Imaging data acquisition and preprocessing were identical to Studies 1-3. The first-level GLM model included five separate boxcar regressors time-logged to the anticipatory cue period corresponding to each rating, which allowed us to model brain activity in response to each trial (i.e., preparatory motor response) separately. The SUITAS was applied to the first-level activation map for each number rating to obtain the signature responses, which were next correlated with the true ratings across participants.

Study 4: visual threat-conditioning dataset

The visual threat-conditioning dataset was from a previous pharmacological study that examined the effect of angiotensin II type 1 receptor antagonist losartan (LT) in threat extinction learning[1]. The brain activations during threat acquisition phase were used to test the generalizability of our predictive model (i.e., SUITAS) in predicting uncertain threat anticipation during associative learning across cohorts, paradigms, MRI systems and scanning parameters.

In this study, 59 male participants (losartan, group: $n = 30$, mean ± SD age = 20.50 ± 1.80; Placebo group: $n = 29$, mean ± SD age = 20.86 ± 1.68) underwent an visual threat-conditioning paradigm during fMRI scanning. Importantly the active treatment (losartan) was only administered following the conditioning procedure, avoiding confounding treatment effects in the context of the current study. Two colored squares (red and blue) were used as conditioned stimuli (CSs). One colored square (red) was designated as the CS+ and was pseudorandomly paired with a mild electric shock (unconditioned stimulus, US, 2 ms) on 43% of the trials, whereas the other colored square (blue) was never paired with shock (CS-). All CSs were presented for 4 s, with a 9-12s interstimulus interval (ISI). The acquisition included two runs and each run consisted of 8 unreinforced CS+, 8 CS-, intermixed with 6 CS+ paired with the shock. The participants were instructed to pay attention to the stimuli and find out the relationship between the stimuli and the shocks. The shock was generated by a Biopac stimulator module STM100C and a STIMSOC adapter (Biopac Systems, Inc.) and the intensity level was determined by a work-up procedure (details and the extinction phase please see ref.[1]).

MRI data were collected on a 3.0-T Siemens TRIO system with a 12-channel head coil (Siemens, Erlangen, Germany). Imaging data were preprocessed and analyzed using SPM12 (scanning parameters, preprocessing and analysis steps see ref.[1]).

Study 5: auditory threat-conditioning dataset

The auditory threat-conditioning dataset was from a previous study that developed a threat-predictive pattern[2]. This dataset served the same purpose as Study 4 with a different modality of conditioned stimuli (CSs).

In this study, 68 participants (45 females, mean ± SD age = 29.64 ± 15.89) underwent an auditory threat-conditioning paradigm during fMRI acquisition. Two different tones (800 and 170 Hz) were used as CSs. One of the tones was designated as the CS+ and was paired with a mild electric shock (unconditioned stimulus, US) on 33% of the trials, whereas the other tone was never paired with shock (CS-). All CSs were presented for 4 s, with a 10 s fixed inter-trial-interval (ITI). Acquisition included 8 randomized repetitions of CS+ paired with shock, 8 randomized repetitions of unreinforced CS+, and 8 randomized repetitions of CS-. The participants were told that they might receive shocks during the experiment without knowing the contingency. The tones were delivered through MR-designed headphones, and were outside of the range of scanner noise. Mild electric shocks were delivered through a bar electrode attached to the right wrist. The shock intensity was determined by a work-up procedure (details and the extinction phase please see ref.[2]).

MRI data were collected on a 3.0-T Siemens Allegra scanner and a 32 channel Siemens head coil. Imaging data were preprocessed and analyzed using SPM8 and custom Matlab (MATLAB, The MathWorks, Inc., Natick, MA) code (scanning parameters, preprocessing and analysis steps see ref.[2]).

Study 6: thermal pain dataset

In the corresponding study by Wager et al.[3] (see also Woo et al.[4]), 33 healthy participants (22 females; mean ± SD age = 27.9 ± 9.0 years) underwent a thermal pain paradigm in which six distinct temperatures (44.3–49.3°C in 1°C increments) were delivered to the left forearm during fMRI acquisition using a TSA-II Neurosensory Analyzer (Medoc Ltd.) with a 16-mm Peltier thermode end-plate. Each thermal stimulation lasted 12.5 s (3 s ramp-up, 2 s ramp-down, and 7.5 s target temperature). After a 4.5–8.5 s jitter, participants judged whether the stimulus was painful or non-painful (4 s) and provided a pain rating on a 100-point visual analogue scale from "no pain" to "worst imaginable pain" (7 s). There are 9 functional runs in total. Runs 1, 2, 4, 8 and 9 include 11 stimulations for each temperature level 1-5. In runs 5-6, temperatures were increased one degree, with 4 stimuli at each of levels 2-6. During run 3 and 7, participants were asked to cognitively regulate the pain intensity in which ten randomly stimulations were presented. The "regulation" runs (run 3 and 7) were not included in generating the thermal pain dataset. Stimulus presentation and behavioral data acquisition were controlled using E-Prime software (PST Inc.).

Whole-brain fMRI data were acquired on a 3T Philips Achieva TX scanner (imaging data acquisition parameters see Woo et al., 2015[4]). Structural and functional MRI data were preprocessed and analyzed using SPM8 (scanning parameters, preprocessing and analysis steps see Woo et al., 2015[4]).

Study 7: monetary gain/loss anticipation dataset

The dataset in Study 7 was taken from an ongoing study from College Student

Cohort of Zhangjiang International Brain Biobank (https://zib.fudan.edu.cn) which used the monetary incentive delay (MID) task, one of the classical and widely used fMRI paradigms for reward processing[5], to assess reward/punishment processing with gain/loss conditions for small or large amounts of money (i.e., -5.0 ¥, -0.2 ¥, 0, 0.2 ¥ or 5.0 ¥) and a neutral (no gain or loss) condition[6]. The present MID task is identical to the MID paradigm from the Adolescent Brain Cognitive Development (ABCD) study[7], and was implemented in healthy young adults in China. In each trial, participants first saw one of three cue shapes (circle, square, or triangle) representing reward, punishment, or no reward/punishment condition, respectively, with the corresponding word inside the cue shape and the amount of money below the cue shape. The cue appears for 2 seconds followed by a 1.5-4 second delay phase (black cross). Participants anticipated reward or punishment during the cue presentation and delay phases. Then, a blank target cue (same shape as the cue presented earlier) appeared on the screen, and participants had to quickly press the response button to either receive a reward or avoid a punishment. After a short delay (1.5-1.85 s), the feedback for the current trial (i.e., the amount of monetary gain or loss) and the accumulated reward will appear. The entire fMRI experiment consisted of 2 runs, each with 50 trials (10 trials per experimental condition), and lasting approximately 5.5 minutes. The study conforms to Fudan University Institutional Review Board's rules and procedures, and all participants provide informed consent. We randomly selected 100 participants (18.68 ± 0.87 years old, 63 females) to control for potential explanation of general anticipation. Stimulus were presented using E-prime Psychology Software (PST Inc.).

Magnetic resonance imaging (MRI) data were collected on a 3T Siemens Prisma. 3-dimensinal T1-weighted images (0.8 mm isotropic, TR = 2500 ms, TE = 2.25 ms) were acquired with a gradient-echo sequence for anatomical localization and co-registration. High spatial (2.0 mm isotropic) and temporal (TR = 800 ms) resolution functional MRI time series were acquired with echo-planar imaging sequence. All functional images were preprocessed with recommended protocols from fMRIPrep (version 20.2.3, https://fmriprep.org)[8] and included the following major steps: (i) T1w-reference workflow; (ii) correction for susceptibility distortions based on fieldmap; (iii) head-motion correction; (iv) co-registration to T1w-reference and normalization to MNI standard space; (v) spatial smoothing with a 6 mm full-width at half-maximum (FWHM) Gaussian kernel and temporal detrending. The first-level GLM model included the combinations of the following regressors of interest Target (Hit or Miss) * Phases (Anticipation, Feedback) * Task Conditions (large-loss, small-loss, neutral, small-win or large-win) and 26 additional covariate regressors (i.e., 24 motion-related parameters: 6 rigid-body motion parameters, their first temporal derivatives and 12 corresponding squared items; as well as mean signals of both white matter and ventricles). In line with the aim of the present study, the contrasts of interest included positive anticipation (mean of small- and large- win anticipation versus implicit baseline) and negative anticipation (mean of small- and large- loss anticipation versus implicit baseline) which were classified by the SUITAS relative to neutral anticipation (no win or loss).

Study 8: visually-induced emotional arousal dataset

Study 6 served to control for arousal and presented arousing emotional and non-arousing neutral pictures during fMRI. The participants ($n$ = 48, 25 females, mean ± SD age = 20.10 ± 2.20 years) of Study 8 were from Study 3 to study emotional processing in healthy adults, in which data from 2 participants were missing. Thirty-two emotionally arousing pictures (16 negative, i.e., disgust, 16 positive, i.e., joy, adoration, amusement) and 16 neutral were selected from the IAPS (International Affective Picture System) and NAPS (Nencki Affective Picture System) and determined by arousal ratings (mean in the 1-7 arousal scale: high-arousing negative, 5.17; neutral, 2.02; high- arousing negative versus neutral: $t_{21} = 10.73$, $P < 0.001$) from an independent sample ($n$ = 22, 8 females, mean ± SD age = 18.95 ± 0.82 years). The participants were asked to passively view the pictures. Stimuli were presented using the E-Prime software (Version 2.0; Psychology Software Tools, Sharpsburg, PA). Stimuli were presented in a single fMRI run (4 min 48 s in total, 2 s picture presentation and 3-5 s fixation per trial) in a pseudorandom order with no more than two consecutive trials of the same category.

Imaging data acquisition and preprocessing were identical to Study 3. The first-level GLM model included one regressor of interest for each experimental condition – the picture viewing period. The SUITAS was tested by means of classifying highly arousing disgust picture viewing period versus implicit baseline (fixation) compared to neutral picture viewing period versus implicit baseline.

<u>Study 9: physiological arousal dataset</u>

We collected skin conductance data during fMRI experiments via an MRI-compatible Biopac system (MP-150) in Studies 1–3. Skin conductance (1000 Hz) was sampled using MRI-compatible disposable, radiotranslucent, pre-gelled electrodes (EL508) attached to the index and middle fingers of the non-dominant (left) hand.

Skin conductance data were downsampled to 100 Hz, filtered using a 1 Hz low-pass filter, and square root-transformed to normalize the distribution using Biopac Acqknowledge 4.2.0 software. Using in-house MATLAB code, we extracted the maximum of the SCL values in a time window of 1-8 s following the cue/anticipation onset and removed the baseline SCL value (the mean) in a 1 s window before the cue onset from the maximum for each trial of each participant, in line with previous methodologies[9]. For the subsequent model prediction, the trials were grouped into quintiles based on the SCL values for each participant and the brain activation maps were binned (i.e., averaged) from 1 to 5 (anticipation period, one map per level) to reflect different SCLs.

Study 10: fear dataset

The fear dataset was from our previous study that developed a subjective fear decoder[10]. The fear induction paradigm used fearful pictures selected from the IAPS (International Affective Picture System), NAPS (Nencki Affective Picture System) and additional pictures from the internet. The study included n = 67 adults (discovery cohort: 34 females, mean ± SD age = 21.5 ± 2.1 years). 80 photographs were distributed in 4 fMRI runs with each presented once. Each trial consisted of a 6-s picture presentation period followed by a 2 s fixation-cross separating the stimuli from the rating period (4 s). Participants reported the fearful state they experienced during the picture presentation using a 5-point Likert scale from 1 (neutral/very slight fear) to 5 (very high fear). Stimuli were presented using E-Prime software (Version 2.0; Psychology Software Tools, Sharpsburg, PA). Another validation cohort in this study includes 20 participants (6 females; mean ± SD age = 21.75 ± 2.61 years) who underwent a similar task in the fMRI scanner as the discovery cohort and also provided ratings of fear experience (details see ref.[10]). The generalization cohort for the fear model include 31 participants (15 females; mean ± SD age = 23.29 ± 4.21 years) who underwent a fMRI session where they were presented with 3600 fearful images consisting of 30 animal categories and 10 object categories (details see ref.[9,10]).

MRI data were collected on a 3.0-T GE Discovery MR750 system (General Electric Medical System, Milwaukee, WI, USA). Structural and functional MRI data were preprocessed and analyzed using SPM12 (scanning parameters, preprocessing and analysis steps see ref.[10]).

Study 11: negative affect dataset

The negative affect dataset was from a previous study developing a multivariate pattern that predicts subjective ratings of negative emotion[11]. A total of 183 participants (female = 52%, mean ± SD age = 42.77 ± 7.3 years) were recruited and divided into a training set (*n* = 121) and a hold-out test dataset (*n* = 61). Stimuli consisted of 15 negative photographs and 15 neutral photographs selected from the IAPS. Each trial begins with a fixation cross (~2 s) followed by a text instruction cue (‘Look’, 2 s). Participants then see a 7-s presentation of negative or neutral images and are asked to rate their emotional state on a Likert scale from 1 (neutral) to 5 (strongly negative).

Finally, there was a jittered rest period (1–3 s). Stimuli were presented using the E-Prime software (Psychology Software Tools, Sharpsburg, PA).

Imaging data were acquired on a 3T Trio TIM whole-body scanner (Siemens, Erlangen, Germany) using a 12-channel, phased-array head coil. fMRI data were preprocessed and analyzed using SPM8 and custom MATLAB (MATLAB, The MathWorks, Inc., Natick, MA) code (scanning parameters, preprocessing and analysis steps see ref.[11]).

### Supplementary Results

<u>Validation of the candidate mechanism of the paradigm – associations between trait anxiety and intolerance to uncertainty and subjective anxious arousal under uncertainty during the UVTA</u>

We ran two separate linear mixed-effects models (LMMs) with trait anxiety (TA) or intolerance of uncertainty (IOU) and uncertainty condition (with safe as baseline: high vs. safe, medium vs. safe, low vs. safe) as fixed effects. With all participants across Study 1–3 ($n$ = 124), we found that there was no significant main effect of TA ($F_{(1,122)}$ = 0.77, $P$ = 0.38), nor interaction between TA score and uncertainty condition ($F_{(2,244)}$ = 0.34, $P$ = 0.71) on subjective ratings, which may reflect a low specificity of the STAI for measuring anxiety[12]. A main effect of uncertainty condition was observed ($F_{(2,244)}$ = 7.76, $P$ < 0.001, $\eta^2_p$ = 0.06 [0.02, 1.00]). In contrast, in the LMM with IOU as a fixed effect in Study 3, significant main effects of IOU score ($F_{(1,48)}$ = 4.13, $P$ = 0.047, $\eta^2_p$ = 0.08 [0.00, 1.00]) and uncertainty condition ($F_{(2,96)}$ = 4.60, $P$ = 0.012, $\eta^2_p$ = 0.09 [0.01,

1.00]) were revealed, yet no interaction effect was found ($F_{(2,96)} = 0.66$, $P = 0.52$). Supplementary Fig.2 shows the mean subjective anxious arousal rating for each condition as a function of intolerance of uncertainty (IOU) score.

SUITAS prediction of the preparatory motor response

To investigate whether our SUITAS model was also sensitive to predict motor response anticipation corresponding to different colors, we applied the SUITAS to the activation maps modeled for each rating of the control experiment in an independent dataset ($n = 20$). Results showed that the SUITAS could not predict the preparatory/anticipatory motor activity based on the colored cue ($r = 0.072$, $p = 0.476$, see Supplementary Fig. 4). Therefore, we ruled out the possibility that the neural patterns of the SUITAS coded motor response.

SUITAS prediction of physiological arousal

A total of $n = 94$ participants from Study 1–3 ($n = 124$) had complete skin conductance data (training dataset: $n = 29$, 12 females, 22.03 ± 2.74; validation dataset: $n = 22$, 11 females, 22.31 ± 2.08; prospective generalization datasets: $n = 43$, 22 females, 20.16 ± 2.21). With these participants, we found that both the SCL ($F_{(3,91)} = 13.01$, $P < 0.001$, $\eta^2 = 0.3$; low uncertainty vs. safe: $t_{93} = 2.51$, $p = 0.014$, Cohen's d = 0.26; medium uncertainty vs. safe: $t_{93} = 3.42$, $p = 0.001$, Cohen's d = 0.35; high uncertainty vs. safe: $t_{93} = 6.17$, $p < 0.001$, Cohen's d = 0.64; low vs. medium uncertainty: $t_{93} = -0.74$, $p = 0.46$, Cohen's d = 0.08; low vs. high uncertainty: $t_{93} = -4.93$, $p < 0.001$, Cohen's d = 0.51; medium vs. high uncertainty: $t_{93} = -4.34$, $p < 0.001$, Cohen's d = 0.45, Supplementary Fig. 7a) and subjective ratings of anxious arousal ($F_{(3,91)} = 399.94$, $P <$

0.001, $\eta^2 = 0.93$; $t_{93} = 10.36\text{-}34.77$, all $P < 0.001$, Cohen's d = 1.07-3.59, Supplementary Fig. 7b) increased as the uncertainty level of the experiment condition increased. However, the trial-wise SCLs and subjective anxious experience did not correlate with each other at the individual level (mean $r = 0.12$, $p = 0.32$, bootstrap test, Supplementary Fig. 7c), suggesting that the objectively measured physiological responses and self-reported subjective experience is dissociated in representing momentary anxious arousal (see recent discussions in ref.[9,13]). To further validate that the neural basis of subjective anxious arousal is dissociated from that of physiological arousal, the SUITAS was applied to the binned beta images of anticipation stage corresponding to different SCLs and results showed that the SUITAS predicted SCL ($r = 0.216$, $p < 0.001$) with a much smaller effect size than predicting subjective ratings ($r = 0.556$, p < 0.001, difference in effect size: $\Delta r = 0.34$, one-tailed permutation test $P < 0.001$, Supplementary Fig. 7d), demonstrating that the SUITAS was specific to predict subjective experience of anxious arousal during uncertain threat anticipation.

Spatial correlations among thresholded maps of SUITAS, VIFS and PINES

The Pearson correlations decreased after thresholding these signatures at uncorrected $P < 0.01$ (SUITAS versus VIFS: $r = 0.02$; SUITAS versus PINES: $r = 0.03$; VIFS versus PINES: $r = 0.06$, one-tailed permutation test all $P < 0.001$) as well as at uncorrected $P < 0.001$ (SUITAS versus VIFS: $r = -0.002$, one-tailed permutation test $P = 0.232$; SUITAS versus PINES: $r = 0.004$, one-tailed permutation test $P = 0.021$; VIFS versus PINES: $r = -0.009$, one-tailed permutation test $P < 0.001$).

After excluding occipital lobe, the spatial correlations among the SUITAS, VIFS and PINES remained comparable using both unthreshold maps (SUITAS versus VIFS: $r = 0.09$; SUITAS versus PINES: $r = 0.03$; VIFS versus PINES: $r = 0.08$, one-tailed permutation test all $P < 0.001$) and thresholded maps at $P < 0.01$ (SUITAS versus VIFS: $r = 0.04$; SUITAS versus PINES: $r = 0.03$; VIFS versus PINES: $r = 0.06$, one-tailed permutation test all $P < 0.001$) as well as uncorrected $P < 0.001$ (SUITAS versus VIFS: $r = 0.003$, one-tailed permutation test $P = 0.069$; SUITAS versus PINES: $r = 0.003$, one-tailed permutation test $P = 0.084$; VIFS versus PINES: $r = 0.03$, one-tailed permutation test $P < 0.001$).

<u>Comparing the SUITAS, VIFS and PINES excluding occipital lobe</u>

Given that previous studies suggest an involvement of the visual cortex in emotion decoding[14] and the present paradigms were different in the visual engagement, we retrained each model excluding the occipital lobe and then compared their performance on predicting anxious arousal, fear  and negative affect. The prediction performance of the SUITAS excluding occipital lobe on subjective experience of anxious arousal (training set: $r = 0.59$; validation set: $r = 0.60$; generalization set: $r = 0.54$; all $P < 0.001$), fear (training set: $r = 0.30$; validation set: $r = 0.29$; one-tailed permutation test all $P < 0.001$; difference in effect sizes ($r$ values): one-tailed permutation test all $P < 0.001$) and negative affect (training set: $r = 0.23$, one-tailed permutation test $P = 0.014$; validation set: $r = 0.28$; one-tailed permutation test $P < 0.001$; difference in effect size ($r$ values): one-tailed permutation test all $P < 0.001$; Supplementary Fig. 8) remained consistent. Similarly, the VIFS excluding occipital lobe more accurately predicted subjective fear (training set: $r = 0.51$; validation set: $r = 0.57$; all $P < 0.001$) than anxious arousal (training set: $r = 0.45$, permutation test $P < 0.001$; validation set: $r = 0.45$, permutation test $P < 0.001$; independent set: $r = 0.24$, permutation test $P = 0.039$) and negative affect (training set: $r = 0.38$, validation set: $r = 0.31$; permutation test all $P < 0.001$), whereas the PINES excluding occipital lobe more accurately predicted subjective experience of negative affect (training set: $r = 0.61$; validation set: $r = 0.74$; all $P < 0.001$) than fear (training set: $r = 0.30$; validation set: $r = 0.36$; permutation test all $P < 0.001$) or anxious arousal (training set: $r = 0.12$, one-tailed permutation test $P = 0.20$; validation set: $r = 0.36$, permutation test $P < 0.001$; independent set: $r = 0.20$; permutation test $P = 0.04$).

<u>ROI- and network-based predictions</u>

For ROI-based predictions, the prediction-outcome correlations for the training dataset (cross-validated, Supplementary Fig. 9a) are: ACC ($r = 0.37, p < 0.001$), aINS ($r = 0.40, p < 0.001$), BNST ($r = 0.22, p < 0.001$), PAG ($r = 0.13, p = 0.06$) and thalamus ($r = 0.23, p < 0.001$); the prediction-outcome correlations for the validation dataset are: ACC ($r = 0.04, p > 0.05$), aINS ($r = 0.30, p < 0.001$), BNST ($r = 0.31, p < 0.001$), PAG ($r = 0.14$, $p = 0.11$) and thalamus ($r = 0.23$, $p < 0.01$); the prediction-outcome correlations for the generalization dataset are: ACC ($r = 0.21$, $p < 0.005$), aINS ($r = 0.38, p < 0.001$), BNST ($r = 0.15, p < 0.05$), PAG ($r = 0.06, p = 0.37$) and thalamus ($r = 0.20, p < 0.005$).

For network-based predictions, the prediction-outcome correlations for the training dataset (cross-validated) are: anterior salience ($r = 0.31$, $P < 0.001$), posterior salience ($r = 0.38$, $P < 0.001$), visual ($r = 0.37$, $P < 0.001$), somatomotor ($r = 0.49$, $P < 0.001$), dorsal attention ($r$ = 0.41, $P < 0.001$), ventral attention ($r$ = 0.45, $P < 0.001$), limbic ($r$ = 0.21, $P < 0.001$), frontoparietal ($r$ = 0.43, $P < 0.001$), default ($r$ = 0.42, $P < 0.001$), and consciousness network ($r$ = 0.54, $P < 0.001$). The prediction-outcome correlations for validation and generalization datasets with different numbers of features tested are shown in Supplementary Fig. 9b.

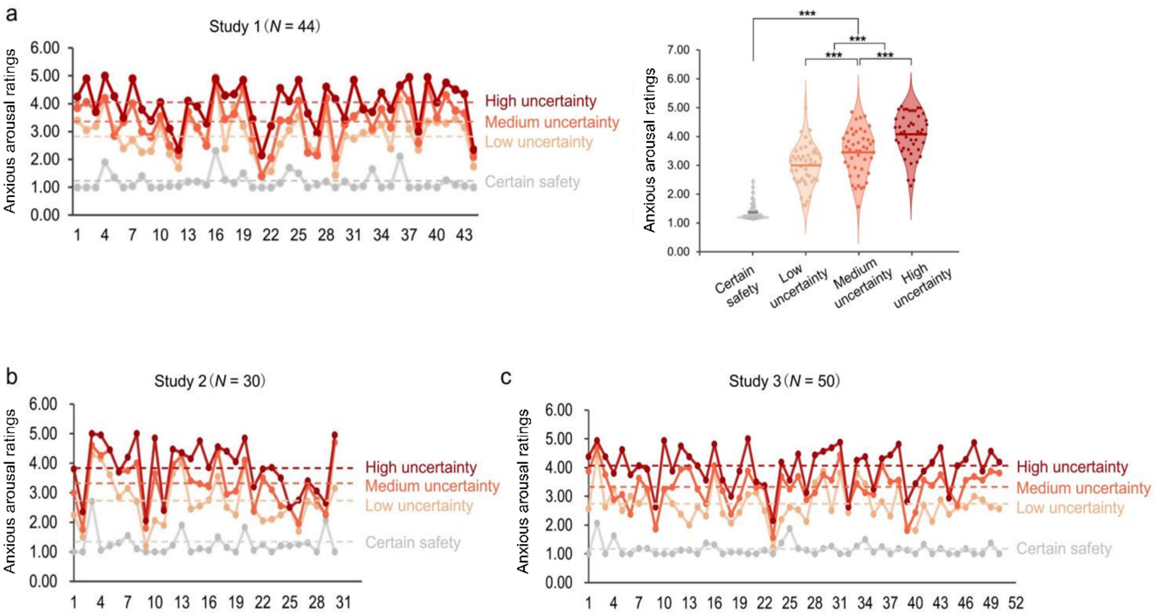


**Supplementary Fig. 1. Effect of shock uncertainty.** The UVTA paradigm induced increased subjective levels of anxious arousal by modulating the uncertainty of outcome. (**a**) anxious arousal ratings for each condition on the individual level (left) and group level (right) in Study 1, (**b**,**c**) anxious arousal ratings for each condition on the individual level in Study 2 and Study 3. The violin plot shows the distribution of anxious arousal ratings for each condition in which the dots represent ratings for each participant and the horizontal lines depict mean values across participants.

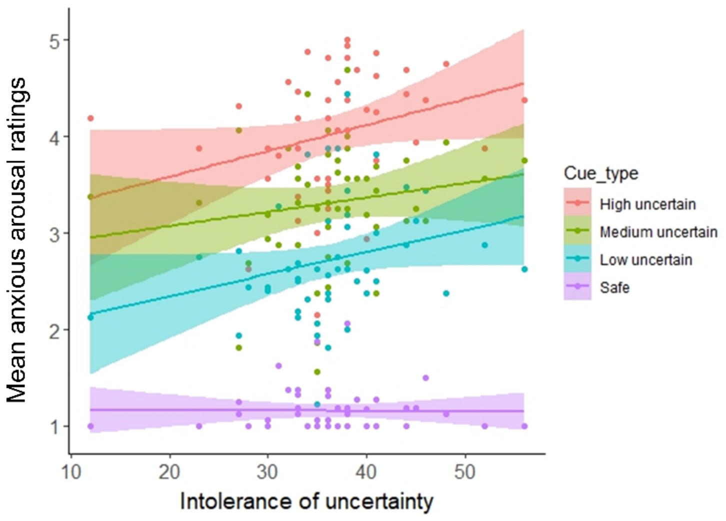


**Supplementary Fig. 2.** Mean subjective anxious arousal rating for each condition as a function of intolerance of uncertainty (IOU) score. Please note that IOU was only assessed in the generalization dataset (Study 3, $n = 50$). Each dot corresponds to a single participant's mean anxious arousal rating and IOU score in one condition. Dashed lines show the linear fit to the data. Shaded areas indicate s.e.

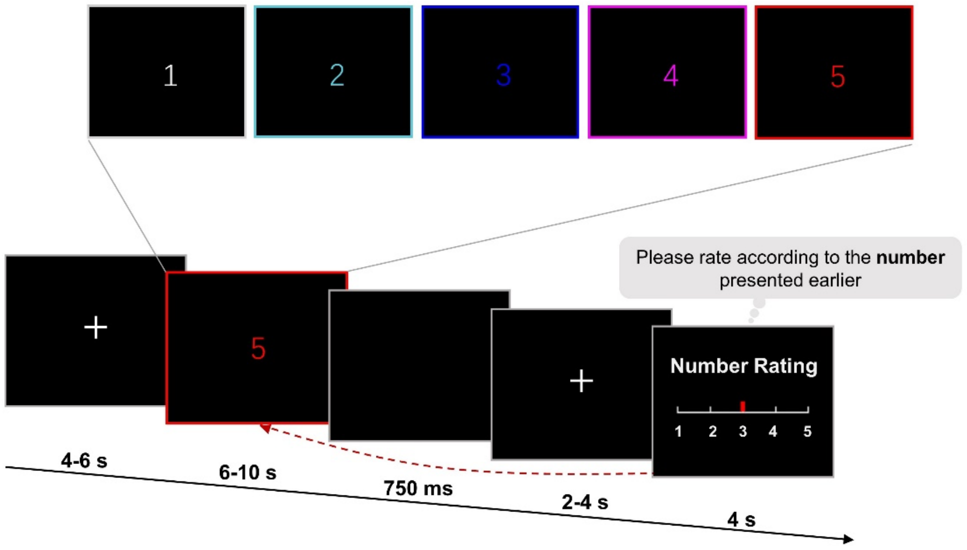


**Supplementary Fig. 3. Experimental design to control for the color- and motor-related effect.** The participants were shown five different colored numbers from 1-5, each with a specific color. The participants were required to anticipate the number that they were going to rate (i.e., preparatory motor response) when the colored number disappeared on a scale ranging from 1 to 5. The duration for each phase was kept the same as the UVTA paradigm.

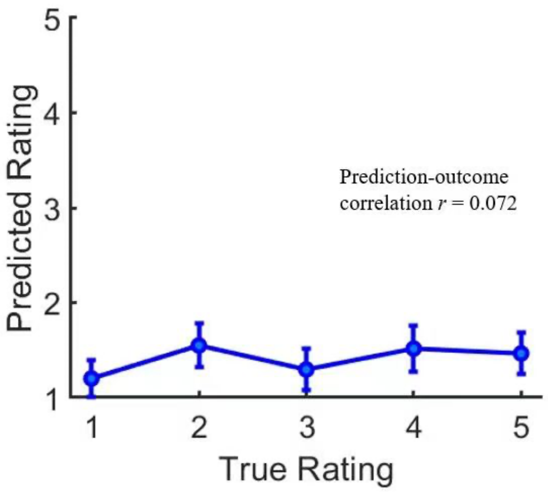


**Supplementary Fig. 4. Using the SUITAS to predict the preparatory motor response corresponding to each of the five colors/numbers.** Predicted rating (signature response) modeled using the anticipation period of the control experiment compared to true ratings across participants in an independent sample (*n* = 20).

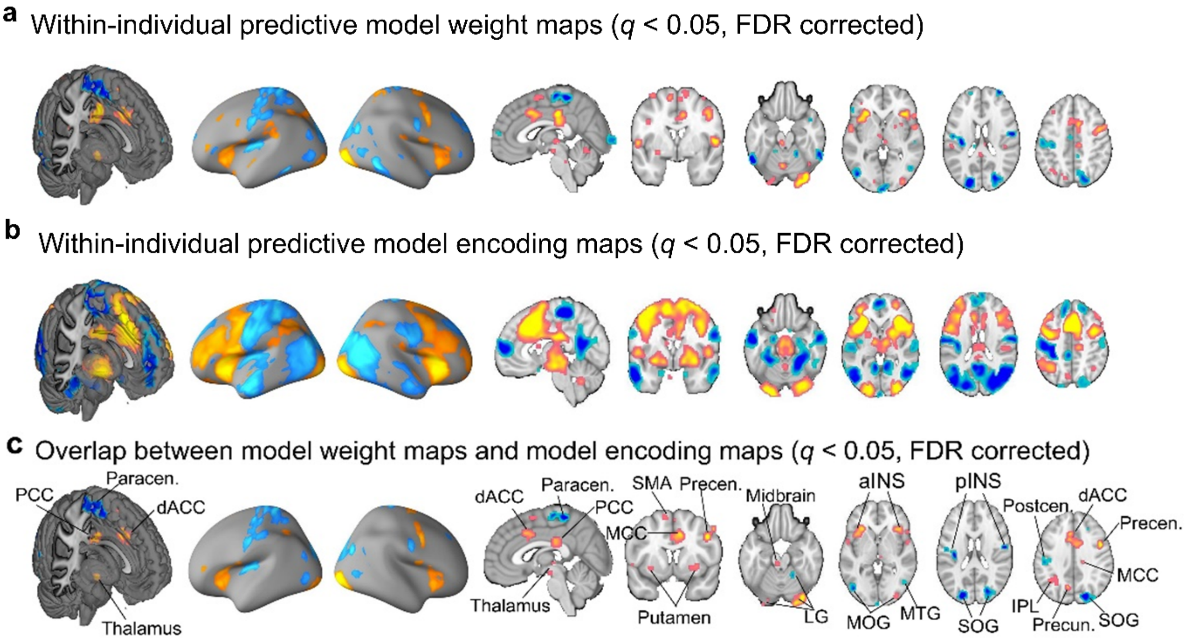


**Supplementary Fig. 5. Within-individual model weight maps, model encoding maps, the overlap and the comparison with population-level model SUITAS.** (**a**) Within-individual model weight maps show regions that consistently predict subjective anxious arousal across participant using a one-sample *t* test (FDR $q < 0.05$) based on training a separate model for each participant's trial-by-trial ratings. (**b**) Within-individual encoding maps show regions that consistently encode subjective experience of anxious arousal and were transformed from within-individual model weight maps using one-sample *t* test (FDR $q < 0.05$). (**c**) The conjunction of within-individual weight maps and encoding maps (FDR $q < 0.05$). Hot color indicates positive values and cold color indicates negative values.

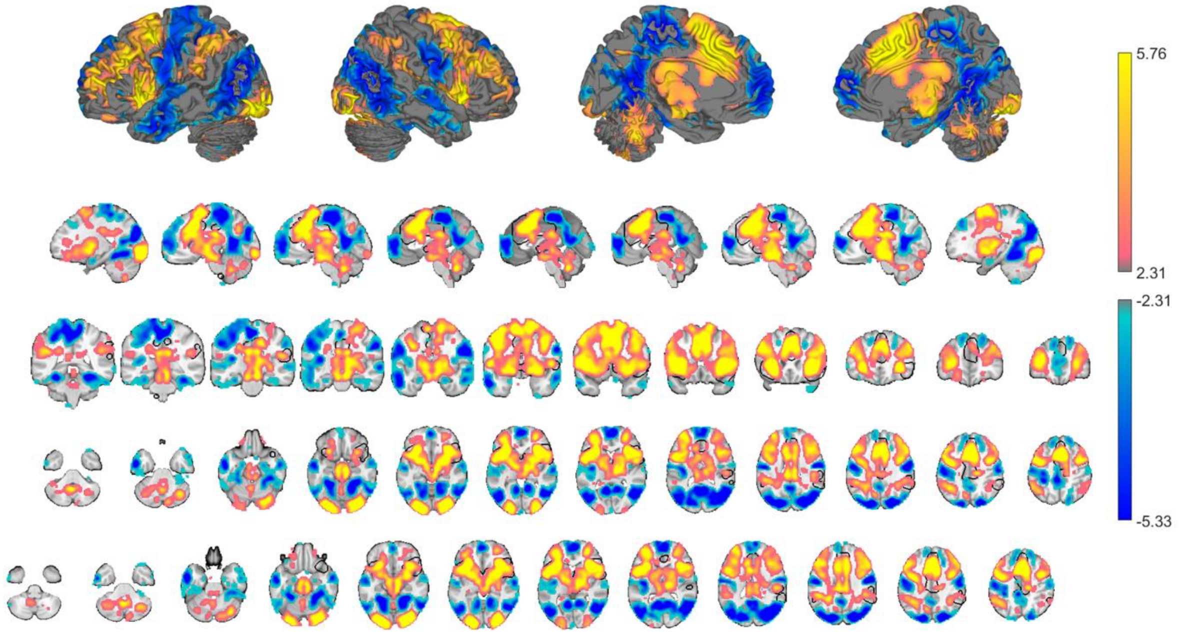


**Supplementary Fig. 6. Univariate categorical results of uncertain threat versus certain safety anticipation.** The univariate activation maps for the contrast of uncertain threat versus certain safety anticipation using the training dataset (FDR $q < 0.05$). Hot color indicates positive values and cold color indicates negative values. The black contour line delineates regions from a meta-analytic activity study of induced anxiety comparing uncertain threat versus safe anticipation in healthy individuals[15].

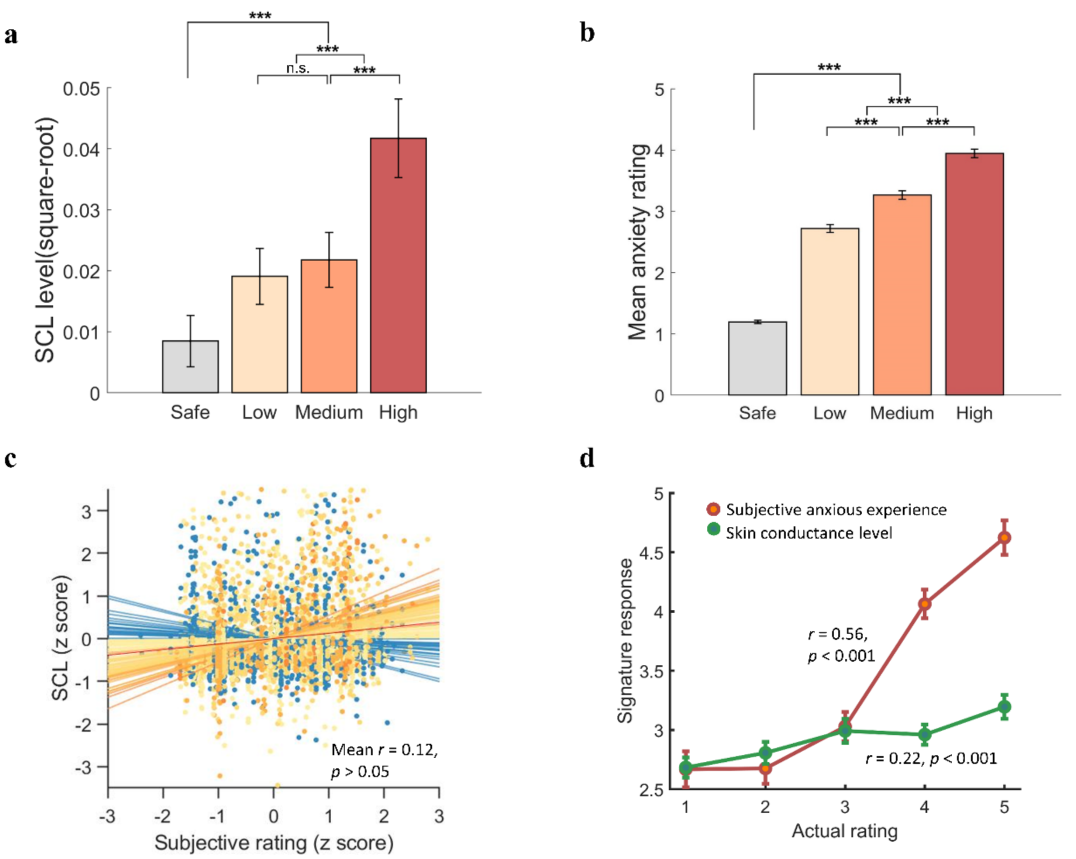


**Supplementary Fig. 7. SUITAS predictions of the subjective experience of anxious arousal and skin conductance levels.** (**a**) The skin conductance levels (square-root) and **(b)** the subjective anxious arousal ratings across participants from training, validation and prospective generalization datasets ($n$ = 94 in total) increased with the uncertainty levels. (**c**) The trial-wise SCL and anxious arousal did not correlate with each other at the individual level. (**d**) SUITAS specifically predicts subjective anxious arousal ratings more than twice than its physiological correlates in terms of the effect size.

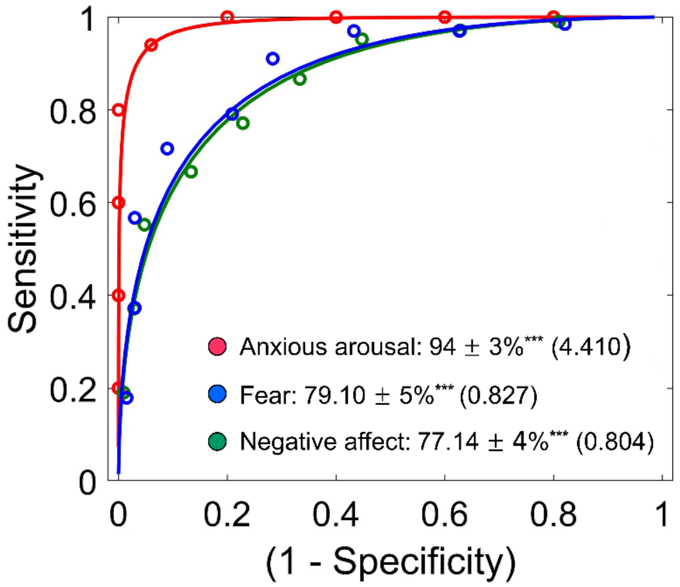


**Supplementary Fig. 8. SUITAS predictions of high versus low subjective anxious arousal, fear and negative affect.** The SUITAS predicted high versus low subjective anxious arousal ratings (prospective generalization dataset, $n$ = 50, Study 3) more accurately compared to high versus low subjective fear in Study 10 ($n$ = 67) and subjective negative affect in Study 11 ($n$ = 121). Performance was shown as accuracy ± SE (Cohen's d). ***$P < 0.001$.

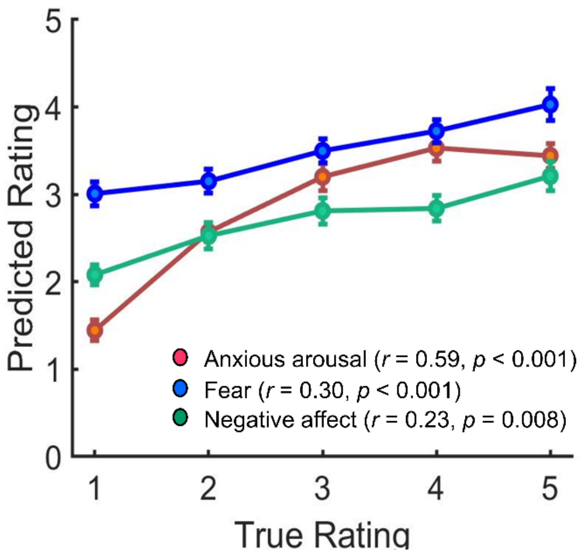


**Supplementary Fig. 9. SUITAS predictions excluding occipital lobe of subjective experience of anxious arousal, fear and negative affect.** The SUITAS predicted ratings compared to the actual ratings for the cross-validated participants in Study 1 (*n* = 44), training dataset in Study 10 (*n* = 67) and Study 11 (*n* = 121). P values were based on permutation tests with 5,000 random shuffles.

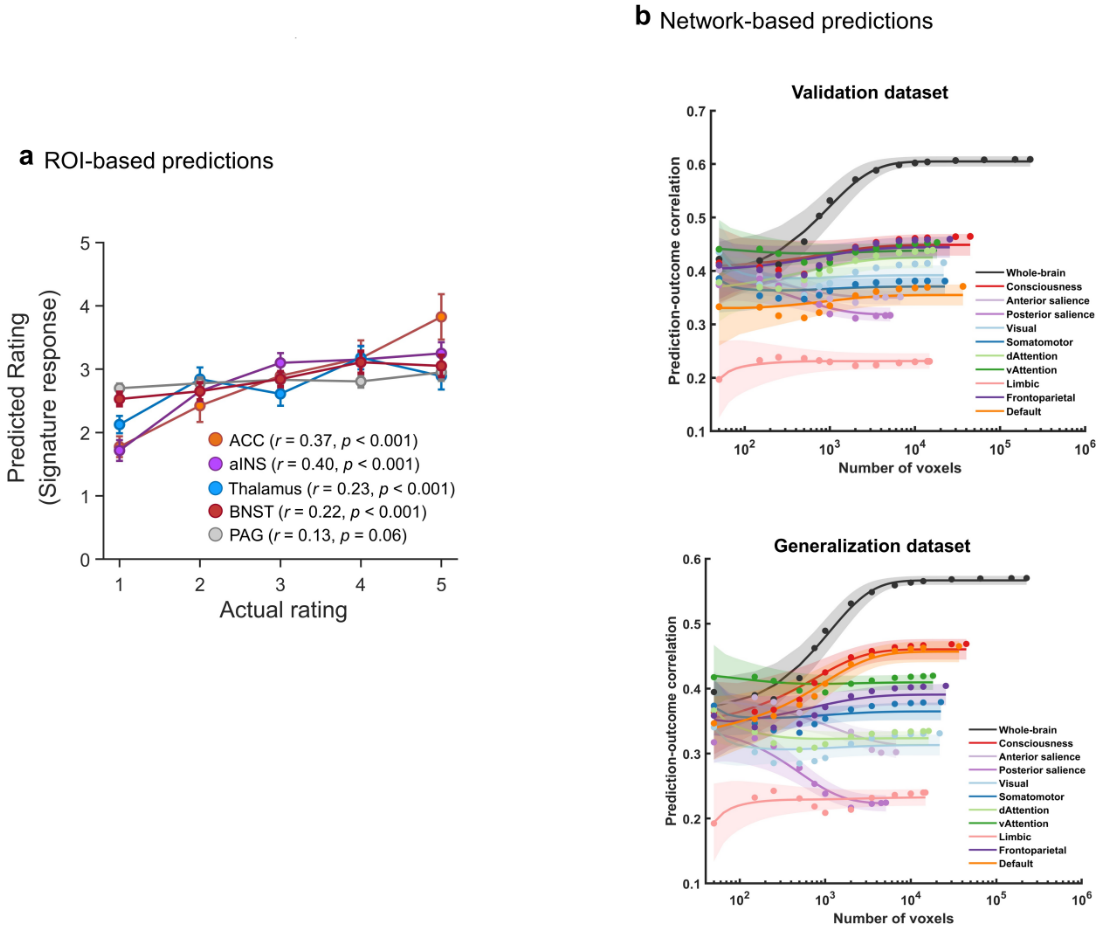


**Supplementary Fig. 10. Local brain region and network predictions.** (**a**) The cross-validated predicted ratings (signature response) compared to the actual ratings in the ROI-based predictions (Study 1). Error bar indicates standard error of the mean. (**b**) Model performance was evaluated as increasing numbers of voxels/features (x axis) to predict subjective anxious arousal in different networks and whole-brain. The models were developed with the training dataset and applied to the validation dataset (top) and the generalization dataset (bottom). Colored dots indicate the mean correlation coefficients, solid lines indicate the mean parametric fit and shaded regions indicate standard deviation. The asymptotic predictions of both validation and generalization datasets when sampling from all brain systems were consistently higher than sampling the same number of voxels from the individual networks when exceeding 1,000 voxels. The consciousness network was composed of anterior cingulate cortex, inferior frontal gyrus, middle frontal gyrus, superior frontal gyrus, orbitofrontal gyrus, rectus, olfactory and insula from the AAL atlas and the posterior parietal cortex from Shirer et al.[16].

**Supplementary Table 1. Experimental parameters for Study 1-11.**

| Study | Dataset | Stimulus type | *N* (female) | # Runs | # Trials / run | Duration for regressor of interest | Rating scale | % Participants rated full scale |
|---|---|---|---|---|---|---|---|---|
| 1 | Liu et al. [a] | electric stimulation | 44 (23) | 4 | 24 | 6-10 s | 1-5 | 59 |
| 2 | Liu et al. [b] | electric stimulation | 30 (14) | 4 | 24 | 6-10 s | 1-5 | 50 |
| 3 | Liu et al. [c] | electric stimulation | 50 (27) | 4 | 29 | 6-10 s | 1-5 | 80 |
| 4 | Zhou et al. 2019 | conditioned visual stimuli. | 59 (0) | 2 | 22 | 4 s | no rating | not applicable |
| 5 | Reddan et al. 2018 | conditioned auditory stimuli | 68 (45) | 4 | 24 | 4 s | no rating | not applicable |
| 6 | Wager et al. 2013 | thermal stimulation | 33 (22) | 7 | 55 for 5 runs, 20 for 2 runs | 12.5 s | 0-100 | 100 (binned 1-6) |
| 7 | ZIB [d] | geometric shapes | 100 (63) | 2 | 50 | 3.5-6 s | no rating | not applicable |
| 8 | Liu et al. [c] | Disgust images (IAPS, NAPS) | 48 (25) | 1 | 32 | 2 s | no rating | not applicable |
| 9 | Liu et al. [b,c] | skin conductance | 65 (33) | 4 | 24/29 | 6-10 s | binned level 1-5 | 100 |
| 10 | Zhou et al. 2021 | fearful images (IAPS, NAPS, internet) | 67 (34) | 4 | 20 | 6 s | 1-5 fear | 97 |
| 11 | Chang et al. 2015 | negative images (IAPS) | 121 | 1 | 30 | 7 s | 1-5 negative | ~80 |

[a] Training dataset; [b] Validation dataset; [c] Generalization dataset; [d] unpublished data from external source; IAPS, International Affective Picture System; NAPS, Nencki Affective Picture System.

**Supplementary Table 2. Prediction performance (correlation) of the VIFS and PINES on anxious arousal, fear and negative affect ratings.**

| Model | Prediction | Anxious arousal | Fear | Negative affect |
|---|---|---|---|---|
| VIFS | Training dataset | 0.35 [0.23, 0.46]; 0.36 ± 0.08 | 0.57 [0.49, 0.63]; 0.89 ± 0.01[a] | 0.35 [0.28, 0.43]; 0.64 ± 0.03 |
| | Validation dataset | 0.35 [0.21, 0.47]; 0.34 ± 0.11 | 0.59 [0.48, 0.69]; 0.87 ± 0.02 | 0.29 [0.18, 0.39]; 0.63 ± 0.04 |
| | Generalization dataset | 0.18 [0.05, 0.31]; 0.28 ± 0.08 | 0.56 [0.45, 0.64]; 0.65 ± 0.06 | |
| PINES | Training dataset | 0.12 [-0.01, 0.24]; 0.20 ± 0.08 | 0.38 [0.28, 0.47]; 0.59 ± 0.04 | 0.60 [0.54, 0.65]; 0.86 ± 0.01[a] |
| | Validation dataset | 0.31 [0.17, 0.45]; 0.38 ± 0.09 | 0.37 [0.21, 0.51]; 0.61 ± 0.07 | 0.72 [0.65, 0.77]; 0.90 ± 0.01 |
| | Generalization dataset | 0.17 [0.03, 0.31]; 0.33 ± 0.08 | 0.20 [0.02, 0.36]; 0.05 ± 0.13 | |

We applied the VIFS and PINES to subjective anxious arousal, fear and negative affect datasets and calculated the overall (bootstrapped 95% CI) as well as within-individual (mean ± SE) prediction-outcome correlations between the signature responses and the actual ratings.

[a] indicates cross-validated

**Supplementary Table 3. Selected ROIs for Fig. 5c and for ROI-based predictions in Supplementary Fig. 10a.**

| ROI | **Areas included** | **Atlas** |
|---|---|---|
| Midcingulate, MCC[a] (supracallosal ACC) | a32pr, a24pr, 33pr, p32pr, p24pr | HCP-MMP v1.0, Glasser et al. 2016[17] |
| ACC[b] | - | Automated Anatomical Labeling (AAL) atlas[18] |
| Anterior insula[a,b] | FOP4, FOP5, AVI, AAIC | HCP-MMP v1.0, Glasser et al. 2016[17] |
| Amygdala[a] | CM, LB, SF | SPM Anatomy Toolbox v2.2c, Eickhoff et al. 2005[19] |
| Thalamus[a,b] | MD, IL, VPL, MGN, LGN, Pulv | Morel 1997[20]; Krauth, 2010[21] |
| PAG[a,b] | dmPAG, vlPAG_L, lPAG_L, vlPAG_R, lPAG_R | Kragel, 2019[22] |
| vmPFC[a] | 10r, 10v, 10d, 10pp, OFC | HCP-MMP v1.0, Glasser et al. 2016[17] |
| dmPFC[a] | 8BM, 8BL, 9m | HCP-MMP v1.0, Glasser et al. 2016[17] |
| vlPFC[a] | 44,45 | HCP-MMP v1.0, Glasser et al. 2016[17] |
| dlPFC[a] | 8Av, p9-46v, 46 | HCP-MMP v1.0, Glasser et al. 2016[17] |
| BNST[b] | BNST | 3T probability mask, Blackford et al., 2017[23] |

[a] indicates labels used in Fig. 6c

[b] indicates labels used in Supplementary Fig. 9a
